# Mitochondrial dysfunction promotes alternative gasdermin D-mediated inflammatory cell death and susceptibility to infection

**DOI:** 10.1101/2021.11.18.469014

**Authors:** Chi G. Weindel, Xiao Zhao, Eduardo Martinez, Samantha L. Bell, Krystal J. Vail, Aja K. Coleman, Jordyn J. VanPortfliet, Baoyu Zhao, Cory J. Mabry, Pingwei Li, A. Phillip West, Jason Karpac, Kristin L. Patrick, Robert O. Watson

## Abstract

Human mutations in mitochondrial-associated genes are associated with inflammatory diseases and susceptibility to infection. However, their mechanistic contributions to immune outcomes remain ill-defined. We discovered that the disease-associated gain-of-function allele *Lrrk2^G2019S^* (leucine-rich repeat kinase 2) promotes mitochondrial hyper-fission, depolarization, and oxidative stress in macrophages. In the presence of *Lrrk2^G2019S^*-dependent mitochondrial perturbations, AIM2 inflammasome activation promotes more cell death but not more pyroptotic IL-1*b* release. Instead, inflammasome activation in *Lrrk2^G2019S^* macrophages triggers gasdermin D (GSDMD)-mediated mitochondrial pores, driving up ROS-mediated RIPK1/RIPK3/MLKL dependent necroptosis. Consequently, infection of *Lrrk2^G2019S^* mice with *Mycobacterium tuberculosis* elicits hyperinflammation and immunopathology via enhanced neutrophil infiltration. By uncovering that GSDMD promotes non-pyroptotic cell death in *Lrrk2^G2019S^* macrophages, our findings demonstrate that altered mitochondrial function can reprogram cell death modalities to elicit distinct immune outcomes. This provides mechanistic insights into why mutations in *LRRK2* are associated with susceptibility to chronic inflammatory and infectious diseases.

**HIGHLIGHTS:**

- Altered mitochondrial homeostasis reprograms cell death modalities
- GSDMD associates with and depolarizes mitochondrial membranes following AIM2 activation
- GSDMD initiates a shift from pyroptotic to necroptotic cell death in *Lrrk2^G2019S^* macrophages
- *Lrrk2^G2019S^* elicits hyperinflammation and susceptibility to infection in flies and mice

## INTRODUCTION

With the recent renaissance in genomics and metabolomics, we have come to appreciate that altered mitochondrial homeostasis and cellular metabolism are major contributors to human disease. The role of mitochondria in controlling cellular physiology is particularly evident in immune cells, where mitochondria regulate and in some cases trigger, antimicrobial defenses. Consistent with the ancestral bacterial origin of the mitochondrion, when pieces of this organelle leak into the cytosol they are sensed as damage-associated molecular patterns (DAMPs) by innate immune pattern recognition receptors. Mito-DAMPs like cytosolic mtDNA engage DNA sensors like cGAS to activate type I IFN expression^1–3^. Other mito-DAMPs like mitochondrial ROS and cardiolipin activate the NLRP3 or AIM2 inflammasomes to elicit pyroptotic inflammatory cell death^4^. Fission and fusion of the mitochondrial network and mitochondrial energetics are additional regulatory nodes that control cytosolic accumulation of mito-DAMPs^5–7^, inflammasome activation^8^, and macrophage polarization^9–11^. Crosstalk between cell death signaling and effector molecules often occurs on the mitochondrial network, and the decision to commit to cell death via one modality versus another is determined in large part by release of proapoptotic mediators and mito-DAMPs from the mitochondrial outer membrane. Mitochondria exert considerable control over the consequences of programmed cell death by integrating stress and damage signals to trigger either immunologically silent (apoptotic) or highly inflammatory (necroptotic)^12,13^ programmed cell death.

To date, we have learned a great deal about how mitochondria contribute to specific immune outcomes by genetically ablating mitochondrial genes and/or treating cells with inhibitors of mitochondrial function. However, we still know very little about how mitochondrial perturbations drive protective or pathogenic immune responses in the context of actual human disease. Human mutations in genes associated with mitochondrial function have been linked to chronic inflammation^14–19^ and susceptibility to infection with pathogens like *Mycobacterium leprae* and *Mycobacterium tuberculosis* (Mtb)^20–24^, which cause the chronic infections leprosy and tuberculosis, respectively. For example, mitophagy genes (e.g. PARK2, PARL, and PINK1) that encode proteins involved in mitochondrial turnover via ubiquitin-mediated selective autophagy have been repeatedly linked to mycobacterial infection risk^23,25–27^. Factors involved in maintaining mitochondrial network integrity like OPA1 and MFN2 are also associated with mycobacterial susceptibility in human GWAS and in studies of macrophages *ex vivo*^24,28–30^. Although these findings and others strongly suggest that mitochondrial proteins are critical players in peripheral immune responses, we know very little about the biology that drives these connections.

One notable mitochondrial-associated gene with numerous poorly understood connections to inflammation and immunity is leucine-rich repeat kinase 2, or LRRK2. LRRK2 is a large, multiple domain-containing protein that functions both as a GTPase and a kinase^31,32^. Because mutations in LRRK2 constitute the greatest known genetic component of familial Parkinson’s disease (PD)^33–35^, much of what is known about LRRK2 function comes from studies of neurons and other cells of the central nervous system (CNS). In CNS cells, LRRK2 has been functionally implicated in neurite regeneration^36,37^, synaptic function^38,39^, autophagy^40–43^, endolysosomal transport^44,45^, and cytoskeleton dynamics^46–48^. Consistent with LRRK2 controlling intracellular membrane trafficking, RAB proteins are thought to be a major group of direct LRRK2 kinase targets^49,50^. LRRK2 phosphorylation of RAB8A has been shown to control endolysosomal homeostasis in macrophages^51,52^. LRRK2 also has many functional connections to mitochondrial homeostasis: it controls mitochondrial Ca2+ flux and opening of the mitochondrial permeability transition pore^53^, it forms a complex with the mitochondrial Rho GTPase Miro to regulate mitochondrial motility and mitophagy kinetics^54,55^, and it influences mitochondrial fission by interacting with DRP1^56–59^.

Despite a number of important mechanistic insights into how LRRK2 functions in a variety of cell types, we still do not understand why LRRK2 is critical in the context of peripheral immune cells. Nor do we know why mutations in LRRK2 upset the immune milieu to trigger Crohn’s disease^16,60–62^ or confer susceptibility to mycobacterial infection^63–65^. Our recent work looking at *Lrrk2^-/-^* macrophages suggests that LRRK2’s role in mitochondrial homeostasis is intimately connected to its ability to control innate immune outcomes^66^. Specifically, we found that ablating LRRK2 leads to mitochondrial destabilization such that mtDNA leaks into the cytosol and chronically activates cGAS; this results in elevated basal levels of *Ifnb1* and interferon stimulated genes (ISGs), rendering macrophages refractory to type I IFN induction when they encounter a pathogen or immune stimulus. These findings motivated us to study the constitutively active *LRRK2^G2019S^* allele as a model to elucidate the molecular mechanisms through which human mitochondrial mutations impact innate immune and infection outcomes. The *LRRK2^G2019S^* mutation is surprisingly prevalent in the human population, and in certain populations (e.g. Ashkenazi Jews and North Africans), the allele accounts for a significant proportion of PD^67–71^. The *LRRK2^G2019S^* allele is also associated with overall increased risk of cancer, especially hormone-related cancer and breast cancer in women^72^. There is growing interest in the *Lrrk2^G2019S^* allele in the context of peripheral infection and inflammation^51,73^. Recent work from Shutinoski et al. reported that mice expressing *Lrrk2^G2019S^* better control *Salmonella enterica* serovar Typhimurium pathogenesis in a sepsis model, and young *Lrrk2^G2019S^* mice infected with reovirus had significantly increased mortality, despite harboring lower viral titers^74^.

Here, we demonstrate that expression of the *Lrrk2^G2019S^* allele causes significant disruption of mitochondrial metabolism in both flies and mice. In murine macrophages, these defects render mitochondria susceptible to enhanced pore formation by the executioner of pyroptosis gasdermin D. This results in reprogramming of cell death pathways (a shift from pyroptosis to necroptosis) and altered release of inflammatory mediators, namely IL1β. Consequently, *Lrrk2^G2019S^* elicits hyperinflammation and susceptibility to Mtb infection in mice and *Pseudomonas entomophila* infection in flies, highlighting an ancient connection between LRRK2’s role in mitochondrial homeostasis and immunity and providing novel insight into the role of LRRK2 mutations in human disease.

## RESULTS

### Mitochondria in *Lrrk2^G2019S^* cells are fragmented and prone to depolarization in response to cellular stress

Previous studies, including work from our lab, reported mitochondrial dysfunction associated with LRRK2 mutations and found that loss of LRRK2 triggers mitochondrial depolarization, network fragmentation, and altered respiratory capacity in macrophages^56,57,66^. To determine how disease-associated human mutations in LRRK2 impact mitochondrial function, we turned to a common allele of LRRK2 that harbors a single amino acid change in the kinase domain (G2019S). This well-studied mutation increases LRRK2 abundance and activity by rendering its kinase domain, which is capable of auto-phosphorylating LRRK2 at several serines, constitutively active^75–79^. We began by qualitatively assessing the structure of the mitochondrial network in wild-type and *Lrrk2^G2019S^* mouse embryonic fibroblasts (MEFs) by immunofluorescence microscopy, staining for the mitochondrial outer membrane protein TOM20. We observed increased fragmentation of *Lrrk2^G2019S^* mitochondria compared to wild-type controls (Fig. 1A). LRRK2-dependent mitochondrial fragmentation has been previously associated with hyperactivation of the mitochondrial fission protein DRP1^56,57,59,80^. Consistent with this, we saw increased levels of the active form of DRP1 p616 relative to total DRP1 in *Lrrk2^G2019S^* bone marrow derived macrophages (BMDMs) by immunoblot (Fig. 1B). While MEFs are ideal for imaging studies due to their large size, we ultimately wanted to assess the structure of the mitochondrial network in *Lrrk2^G2019S^* BMDMs. To achieve this quantitatively, we developed a strategy utilizing flow cytometry to measure mitochondrial size differences down to the sub-micron level. Polystyrene beads of 1, 2, and 4 μm were used to create a gating strategy to identify mitochondria, as previously described in^81,82^ (Fig. 1C and S1A left plot). Crude and purified mitochondrial isolates stained with or without the mitochondrial specific dye MitoTracker green were used to validate this gating strategy, (Fig. S1A middle and right plots). To quantify mitochondrial size differences between wild-type and *Lrrk2^G2019S^* BMDMs, isolated MitoTracker green-stained mitochondria were compared to size standard beads using the forward scatter (FSC) optical detector (Fig. 1D, E). Consistent with DRP1 hyperactivation, we measured more “small” mitochondria (<1 μm) and fewer “large” mitochondria (>4 μm) in *Lrrk2^G2019S^* BMDMs compared to wild-type (Fig. 1F). Importantly, treating macrophages with low concentrations of the DRP1 inhibitor Mdivi-1 (10μM) significantly reduced the *Lrrk2^G2019S^* mitochondrial fragmentation phenotype (Fig. 1D-F), supporting a role for pDRP1 in promoting mitochondrial hyper fission in these cells.

**Figure 1.**
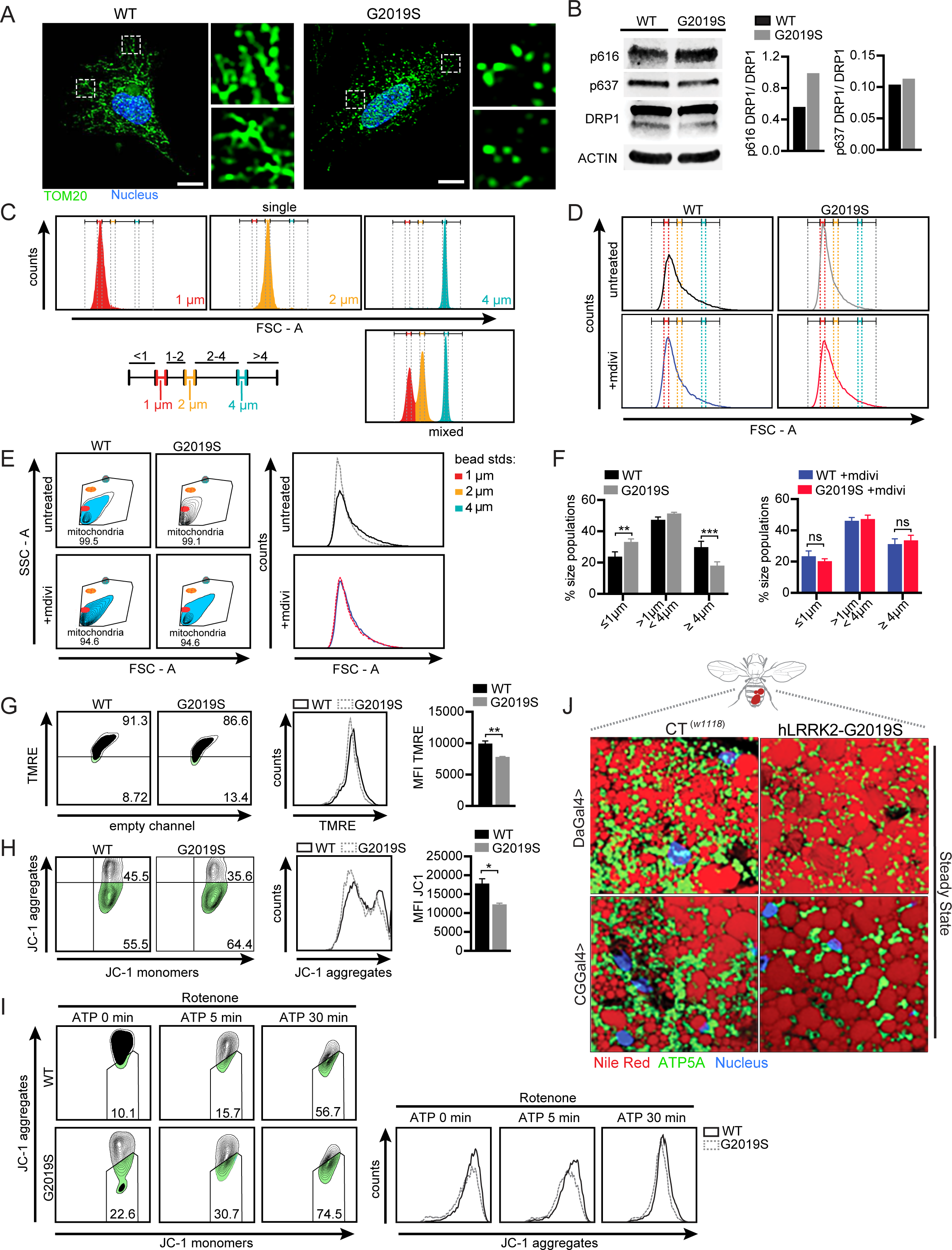
*Lrrk2* G2019S promotes mitochondrial dysfunction marked by network fragmentation and susceptibility to depolarization upon cellular stress. **A.** Immunofluorescence microscopy images visualizing mitochondria (anti-TOM20; green) in wild-type and *Lrrk2^G2019S^* MEFs. Nuclei visualized by DAPI (blue). **B.** Immunoblot analysis and quantification of DRP1 phosphorylation (pSer616 = activation; pSer637 = inhibition) in wild-type and *Lrrk2^G2019S^* BMDMs relative to total DRP1. Beta-actin used as a loading control. **C.** Validation of polystyrene bead size standard resolution at the 1-4μm level. **D.** Flow cytometry of isolated mitochondria from wild-type and *Lrrk2^G2019S^* BMDMs +/-10 µM Mdivi-1 for 16h. **E**. Size resolution of mitochondria from wild-type and *Lrrk2^G2019S^* BMDMs treated with or without Mdivi-1, 10 µM, for 16h. **F.** Quantification of mitochondrial size distribution in wild-type and *Lrrk2^G2019S^* BMDMs based on size standard polystyrene beads +/-10 µM Mdivi-1 **G.** Mitochondrial membrane potential measured by flow cytometry using the cationic dye TMRE (585/15) in wild-type and *Lrrk2^G2019S^* BMDMs. Histograms display (585/15) x-axis. Quantitation shown on the right. **H.** Mitochondrial membrane potential measured by flow cytometry using the cationic carbocyanine dye JC-1. JC-1 Aggregation (610/20) = normal mitochondrial membrane potential; monomers (520/50) = low membrane potential. Histograms display (610/20) x-axis. Quantification shown on the right. **I.** Flow cytometry analysis of JC-1 dye staining in BMDMs treated with rotenone (2.5 µM for 3h) followed by ATP (5 mM for 5 and 30 min.). Right histograms display (610/20) x-axis. **J.** Immunofluorescence microscopy images of *Drosophila melanogaster* fat bodies (Nile Red, red) mitochondria (ATP5A, green), and nuclei (DAPI, blue) in wild-type (CT(w1118)) and mutant hLRRK2-G2019S-expressing flies under a ubiquitous (Da-Gal4) or fat body specific promoter (CGGal4). Data are expressed as a mean of three or more biological replicates with error bars depicting SEM. Statistical significance was determined using a two-tailed Student’s T test: * = p<0.05, ** = p<0.01, *** = p<0.001.

Because mitochondrial fragmentation has been shown to alter membrane potential and responses to stress^83^, we turned to the cell permeable dyes Tetramethylrhodamine (TMRE) and JC-1. We found reduced TMRE fluorescence intensity in *Lrrk2^G2019S^* BMDMs by flow cytometry compared to wild-type cells, indicating mitochondrial depolarization (Fig. 1G). We observed a similar phenotype in cells treated with the dye JC-1, which accumulates as red fluorescent aggregates in healthy mitochondria but forms green fluorescent monomers when membrane potential is lost. We found that *Lrrk2^G2019S^* macrophages had reduced JC-1 aggregates at rest (Fig. 1H) and following treatment with rotenone and ATP (Fig. 1I), which stresses mitochondria by inhibiting complex I. This suggests that expression of the gain-of-function *Lrrk2^G2019S^* allele is sufficient to alter mitochondrial homeostasis in macrophages and sensitize the mitochondrial network to stress.

We recently showed that enhanced mitochondrial dynamics and mobilization of lipid stores are integral to the immune response to bacterial pathogens in the fruit fly *Drosophila melanogaster*^84^. To date, several labs successfully used *Lrrk2^G2019S^*-expressing flies as a model to study LRRK2 function^85–88^. To test whether expression of hLRRK2 influences mitochondrial homeostasis in a manner that is evolutionarily conserved, we utilized transgenic flies expressing hLRRK2-G2019S ubiquitously (DaGal4>) or targeted to the fat body (CGGal4>), an adipose-like tissue that is essential for energy storage (lipid droplets) and innate immunity in insects. The fat body incorporates nutrient and pathogen sensing mechanism in a distinct tissue, before these systems evolved into more complex organ types in mammals^84^. Using immunofluorescence microscopy, we visualized lipid droplets (NileRed) with mitochondria (anti-ATP5A) and observed that the mitochondrial network in the fat body of hLRRK2-G2019S-expressing flies is less dense compared to controls (Fig. 1J, green, right). We also discovered that fat body lipid droplets, which store triacylglycerols, a major fuel source for mitochondrial OXPHOS, are twice as large in hLRRK2-G2019S-expressing flies (Fig. 1J, red). Changes in lipid and/or energy storage, as well as lipid droplet biology, have been linked to aberrant innate immune responses across taxa^84,89,90^. Together, these observations are consistent with *Lrrk2^G2019S^* altering mitochondrial homeostasis and energetics, both in cells *ex vivo* and at an organismal level, in evolutionarily distant animals.

### *Lrrk2^G2019S^* macrophages do not exhibit altered transcriptional responses following innate stimuli

Having observed altered capacity for mitochondrial stress, we next investigated the consequences of altered mitochondrial homeostasis on innate immune responses in *Lrrk2^G2019S^* BMDMs. Many studies have linked mitochondrial function and mito-DAMP release with innate immune outcomes, including type I IFN expression downstream of cGAS-dependent cytosolic surveillance (Fig. 2A, middle), immunologically silent programmed cell death via apoptosis (Fig. 2A, top), and inflammatory programmed cell death via pyroptosis (Fig. 2A, bottom). Consistent with this dogma, our earlier work linked DRP1-dependent mitochondrial fission defects in *Lrrk2^-/-^* macrophages with release of mtDNA into the cytosol, elevated basal type I IFN expression, and an inability to induce a type I IFN response upon infection with pathogens like Mtb^66^. To determine whether *Lrrk2^G2019S^* BMDMs also show altered type I IFN responses, we collected total RNA from wild-type and *Lrrk2^G2019S^* BMDMs at rest and at 4h post-infection with Mtb (MOI = 10), and measured *Ifnb1* and ISGs by RT-qPCR. We have shown in several previous studies that 4h is a key early innate immune time point at which *Ifnb1* expression levels peak in Mtb-infected macrophages^91–93^. Curiously, we observed no differences in *Ifnb1* or ISG expression between wild-type and *Lrrk2^G2019S^* BMDMs, at rest (Fig. S2A), at 4h post-Mtb infection (Fig. 2B), or in response to a variety of other innate immune agonists that stimulate IRF3-dependent gene expression (i.e. dsDNA (ISD), dsRNA (poly I:C), LPS, Pam3CSK4, or CpG (2935) (Fig. S2B-C)). We also observed no significant differences in other proinflammatory transcripts after Mtb infection (*Tnf*, *Il1a*, *Il1b*, and *Il6* shown; Fig. S2D) or stimulation with innate immune agonists (*Tnf* and *Il1b* shown; Figure S2E). From these data, we concluded that *Lrrk2^G2019S^* does not promote chronic engagement of cytosolic nucleic acid sensors, nor does it alter the ability of macrophages to transcriptionally activate inflammatory gene expression following innate immune stimuli.

**Figure 2.**
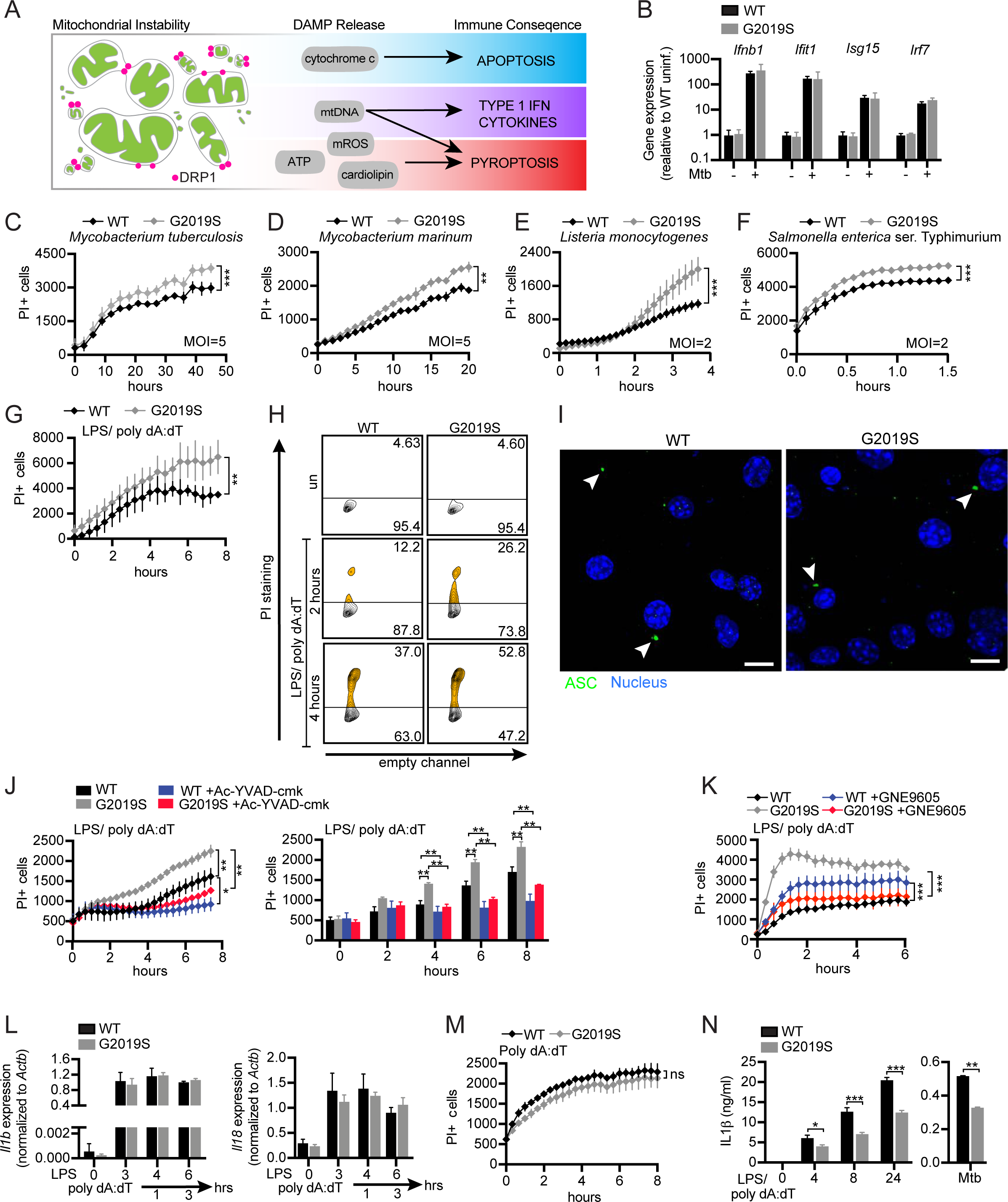
*Lrrk2^G2019S^* promotes cell death during intracellular bacterial infection and AIM2 inflammasome activation. **A.** Model depicting different consequences associated with mitochondrial fragmentation and stress. **B.** Quantification of type I IFN gene expression in wild-type and *Lrrk2^G2019S^* BMDMs by qRT-PCR at 4h post-Mtb infection (MOI = 10). **C.** Propidium iodide (PI) staining over a time course of infection with Mtb (MOI 5). **D.** As in C, but with *Mycobacterium marinum* (MOI = 5). **E.** As in C, but with *Listeria monocytogenes* (MOI = 2). **F.** As in C, but with *Salmonella enterica* (serovar Typhimurium) (MOI = 2). **G.** PI staining of wild-type and *Lrrk2^G2019S^* BMDMs over a time course of AIM2 stimulation. All AIM2 stimulations were performed in BMDMs under the following conditions: 10 ng/ml LPS for 3h followed by 1 µg/ml poly dA:dT. Time courses begin with the introduction of poly dA:dT. **H.** Flow cytometry of PI staining (y-axis) and empty channel (x-axis) in wild-type and *Lrrk2^G2019S^* BMDMs at 2h and 4h post-AIM2 stimulation, as described in G. **I.** Immunofluorescence microscopy images of ASC speck formation during AIM2 activation (anti-ASC, green). Nuclei visualized with DAPI (blue). **J.** As in G but with caspase inhibitor Ac-YVAD-CMK (100 µM). Quantification of PI incorporation at each hour on the right. **K.** As in G but with LRRK2 inhibitor GNE9605 (1 µM). **L.** *Il1b* and *Il18* transcript levels in wild-type and *Lrrk2^G2019S^* BMDMs as measured by qRT-PCR following AIM2 stimulation (1 and 3h) or LPS priming alone (3h). **M.** As in G. but without LPS priming. **N.** Extracellular IL-1*β* protein levels as measured by ELISA over a time course of AIM2 stimulation (as described in G) or at 24h after infection with Mtb (MOI 10). Data are expressed as a mean of three or more biological replicates with error bars depicting SEM. Statistical significance was determined using a two-tailed Student’s T test: * = p<0.05, ** = p<0.01, *** = p<0.001.

### *Lrrk2^G2019S^* macrophages are prone to caspase-1-mediated cell death in response to intracellular bacterial infection and inflammasome activation

Mitochondrial fragmentation and stress can trigger different kinds of regulated cell death (Fig. 2A top and bottom). To determine if *Lrrk2^G2019S^* BMDMs are more or less prone to undergo regulated cell death, we infected wild-type and *Lrrk2^G2019S^* BMDMs with intracellular bacterial pathogens known to activate multiple cell death pathways and measured propidium iodide (PI) incorporation over time using a BioTek Lionheart FX Live Cell Imager: *Mycobacterium tuberculosis* (MOI=5) (Fig. 2C), *Mycobacterium marinum* (MOI=5) (Fig. 2D), *Listeria monocytogenes* (MOI=2) (Fig. 2E), and *Salmonella enterica* serovar Typhimurium (MOI=2) (Fig. 2F). Timescales were adjusted to account for dramatic differences in the dynamics of infection with each of these pathogens (e.g. Mtb replicates in a macrophage monolayer over several days, while *Listeria* will completely overtake a plate of macrophages within 8h). While infection of both wild-type and *Lrrk2^G2019S^* BMDMs led to increased PI incorporation over time, PI levels were 20-100% more in *Lrrk2^G2019S^* BMDMs during these intracellular bacterial infections. We do not believe this phenotype is due to differences in bacteria numbers since the data shown are at timepoints that, for each pathogen, precede substantial intracellular replication. Moreover, we do not observe significant differences in Mtb replication in wild-type and *Lrrk2^G2019S^* BMDMs over a 72h time-course (Fig. S2F). Importantly, we observed no differences in PI incorporation between wild-type and *Lrrk2^G2019S^* BMDMs at rest or in response to stimulation with immune agonists like LPS, Pam3CSK4, or dsDNA (Fig. S2G), suggesting that *Lrrk2^G2019S^* BMDMs are specifically prone to cell death following a complex inflammatory trigger.

Each of these pathogens can elicit multiple types of regulated cell death and PI incorporation does not distinguish between different cell death modalities. Therefore, to begin to understand the nature of the *Lrrk2^G2019S^* macrophage cell death phenotype, we asked whether *Lrrk2^G2019S^* BMDMs were prone to apoptosis, which is triggered downstream of mitochondrial fragmentation and stress leading to cytochrome C release and caspase-3, -7, -9 activation^94^. We observed no differences in annexin V staining between wild-type and *Lrrk2^G2019S^* BMDMs at rest or following treatment with the DNA damage agent etoposide, indicating that *Lrrk2^G2019S^* BMDMs are not more prone to apoptosis (Fig. S2H).

To determine if *Lrrk2^G2019S^* BMDMs are more prone to cell death downstream of inflammasome activation we treated wild-type and *Lrrk2^G2019S^* BMDMs with LPS and the synthetic dsDNA sequence, poly dA:dT, which together are a well-characterized agonist of the AIM2 inflammasome^95^. We observed that AIM2 inflammasome activation, as measured by PI incorporation, was sufficient to recapitulate the enhanced cell death we observed in *Lrrk2^G2019S^* BMDMs during bacterial infection (Fig. 2G). We confirmed this result via flow cytometry measuring PI+ cells at 2h and 4h post LPS/poly dA:dT treatment (Fig. 2H). Consistent with inflammasome activation, we visualized ASC specks via immunofluorescence microscopy but qualitatively observed no significant differences in the percentage of cells containing specks in wild-type versus *Lrrk2^G2019S^* BMDMs at 2h post-AIM2 stimulation (Fig. 2I). Importantly, we found that treatment with Ac-YVAD-cmk, a highly selective and irreversible inhibitor of caspase-1, rescues *Lrrk2^G2019S^*-dependent enhanced cell death triggered by LPS/poly dA:dT (Fig. 2J). Inhibiting LRRK2 kinase activity with the highly potent and selective inhibitor GNE-9605 also rescued the elevated cell death phenotype in a similar fashion (Fig. 2K). Collectively, these data support inflammasome activation as a major driver of *Lrrk2^G2019S^* kinase-dependent cell death during bacterial infection of macrophages.

One possible explanation for *Lrrk2^G2019S^*-dependent inflammasome-triggered cell death is that *Lrrk2^G2019S^* macrophages are “primed” by mito-DAMP release, resulting in transcriptional upregulation of inflammasome components. To test this hypothesis, we measured AIM2 inflammasome-associated transcripts at rest and over a time-course of AIM2 inflammasome activation in wild-type and *Lrrk2^G2019S^* BMDMs. We measured no significant differences in transcript abundance of pyroptotic cell death mediators (*Aim2*, *Nlrp3*, *Pycard*, *Gsdmd* (Fig. S2I)) or inflammasome associated cytokines (*Il18* and *Il1b* (Fig. 2L)). We also saw no elevated cell death after poly dA:dT treatment alone (Fig. 2M), effectively ruling out “priming” as the driver of enhanced cell death in *Lrrk2^G2019S^* macrophages. We then measured IL-1β release as another canonical read-out of AIM2 inflammasome activation. Unexpectedly, while *Lrrk2^G2019S^* BMDMs experience enhanced caspase-1-dependent cell death following AIM2 activation, they release markedly less of the pyroptotic inflammatory mediator IL-1β over a time-course post-AIM2 stimulation (Fig. 2N left) and at 24h post-Mtb infection (Fig. 2N right). Taken together, these data indicate that while inflammasome activation triggers enhanced regulated cell death in *Lrrk2^G2019S^* BMDMs, this death does not follow the paradigm of inflammasome-mediated pyroptosis (i.e. IL-1β release).

### Inflammasome activation triggers additional mitochondrial stresses in *Lrrk2^G2019S^* macrophages

Previous studies have identified connections between mitochondrial stress, mito-DAMP release, and re-programming of cell death signaling pathways^96,97^. Because our data suggest that *Lrrk2^G2019S^* BMDMs undergo non-pyroptotic cell death following treatment with AIM2 agonists, we hypothesized that underlying mitochondrial dysfunction in *Lrrk2^G2019S^* BMDMs skews cell death signaling downstream of inflammasome activation. Thus, we set out to define how mitochondrial homeostasis is altered following inflammasome activation in *Lrrk2^G2019S^* BMDMs.

To begin, we used flow cytometry to measure mitochondrial membrane potential in wild-type and *Lrrk2^G2019S^* BMDMs at 2h post-LPS/poly dA:dT treatment, using FCCP treatment (a potent mitochondrial OXPHOS uncoupler) as a positive control. Compared to wild-type macrophages, resting *Lrrk2^G2019S^* BMDMs showed reduced TMRE fluorescence intensity, which was further reduced by LPS/poly dA:dT treatment (Fig. 3A), consistent with AIM2-dependent mitochondrial depolarization. Next, to determine if mitochondrial depolarization in *Lrrk2^G2019S^* BMDMs promotes mito-DAMP release upon AIM2 activation, we quantified mtDNA in the cytosol of wild-type and *Lrrk2^G2019S^* macrophages. Briefly, we isolated cytosol and membrane fractions via differential centrifugation (as in^66^) and measured ATP5A1 and VDAC1, two abundant mitochondrial proteins, by immunoblot to ensure cytosolic fractions were not contaminated by mitochondrial fragments (Fig. 3B). Subsequent qPCR of the cytosol from wild-type and *Lrrk2^G2019S^* macrophages revealed significantly higher levels of two mtDNA-encoded genes (*16s* and *ND4*) in the cytosol of *Lrrk2^G2019S^* macrophages at 3h post-LPS/polydA:dT treatment (Fig. 3C). Consistent with enhanced cytosolic mtDNA release, we observed higher transcriptional activation of type I IFN expression in *Lrrk2^G2019S^* macrophages compared to wild-type at 4-5h post-AIM2 activation (Fig. S3A). While AIM2 activation is sufficient to release mtDNA in wild-type BMDMs, this release is enhanced in *Lrrk2^G2019S^* mitochondria (Fig. 3C).

**Figure 3.**
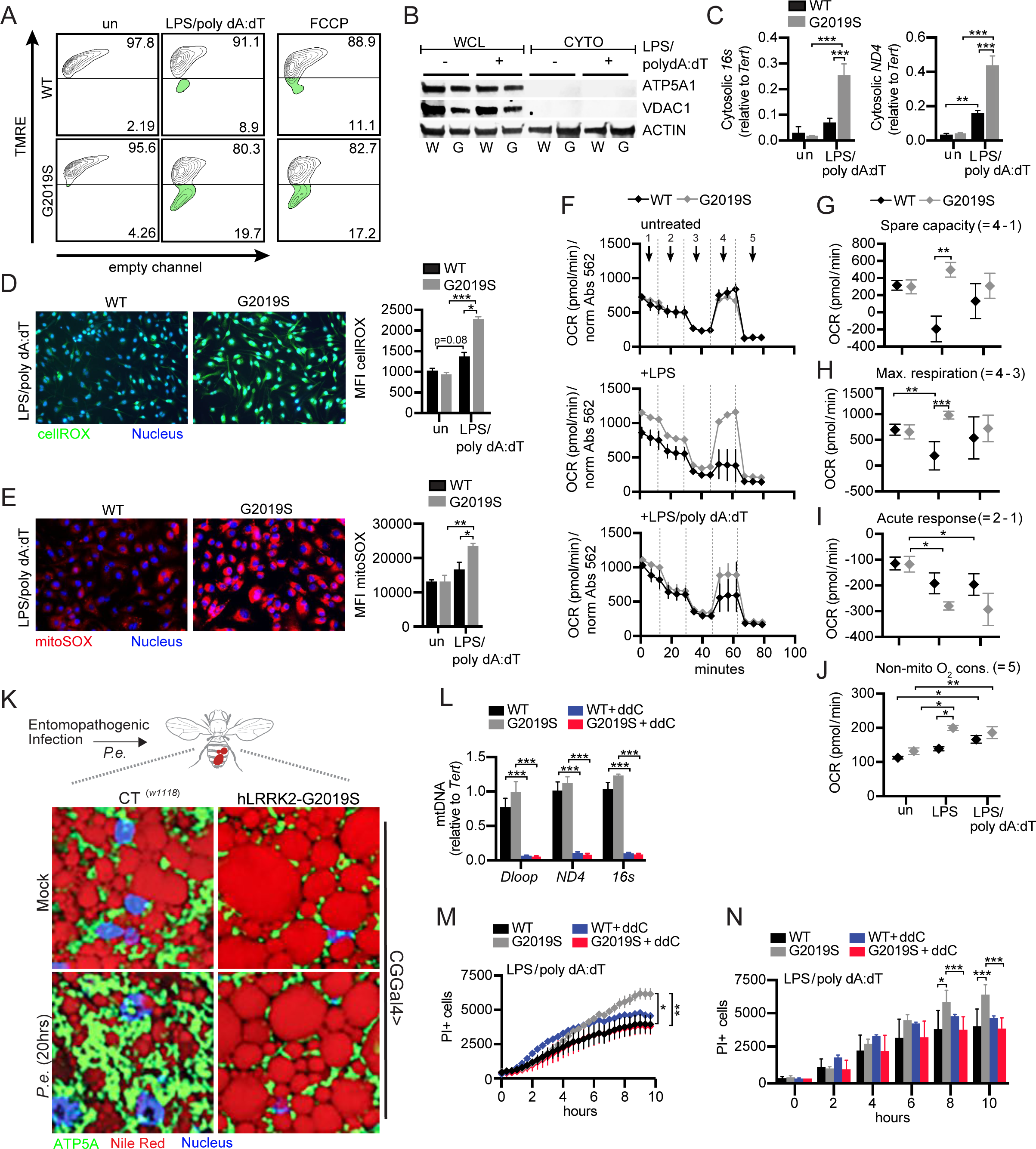
Inflammasome activation triggers additional mitochondrial stresses that alter metabolism and promote cell death in *Lrrk2^G2019S^* macrophages. TMRE staining of wild-type and *Lrrk2^G2019S^* BMDMs 2h post-AIM2 stimulation. 50 µM FCCP treatment (30 min) was used as a positive control. **B**. Immunoblot of total and cytosolic fractions from wild-type and *Lrrk2^G2019S^* BMDMs 3h post-AIM2 stimulation. Mitochondrial proteins ATP5A1 and VDAC1 were used to determine purity of the cytosolic fraction. Beta-actin served as a loading control. **C.** qPCR of mtDNA (*16s* and *ND4*) from cytosolic fractions in B. quantified relative to total nuclear DNA *Tert* in unstimulated and AIM2 stimulated (3h) wild-type and *Lrrk2^G2019S^* BMDMs **D.** Immunofluorescence microscopy images of wild-type and *Lrrk2^G2019S^* BMDMs stained with fluorogenic oxidative stress probe cellROX (green) and live cell nuclear stain NucBlue (blue) 2h post-AIM2 activation. Quantitation of cellROX MFI on right. **E.** As in D but with the mitochondrial targeted superoxide dye mitoSOX. Quantification of mitoSOX MFI on right. **F.** Oxygen consumption rate (OCR) measured by Agilent Seahorse Metabolic Analyzer for wild-type and *Lrrk2^G2019S^* BMDMs: untreated (top), +10 ng/ml LPS 3h (middle), and +10 ng/ml LPS 3h, 1 µg/ml poly dA:dT 3h (bottom). Numbers represent readings after each injection point: Basal reading (no treatment), 2. Glycolysis activation (glucose), 3. ATP synthase inhibition (Oligomycin and sodium pyruvate), 4. Mitochondrial uncoupling (FCCP), 5. Respiratory chain inhibition (Antimycin A and rotenone). **G.** Spare respiratory capacity of untreated, LPS-treated, and AIM2-stimulated wild-type and *Lrrk2^G2019S^* BMDMs. **H**. As in G but measuring maximal respiration. **I.** As in H but measuring acute response. **J.** As in G but measuring non-mitochondrial oxygen consumption. **K**. Immunofluorescence microscopy images of fat bodies (Nile Red, red) mitochondria (ATP5A, green), and nuclei (DAPI, blue); in WT (CGGal4>CT*^(w1118)^*) and hLRRK2-G2019S expressing (CGGal4>hLRRK2-G2019S) *Drosophila melanogaster* infected with *Pseudomonas entomophila* (*Pe*) for 20h. **L.** qPCR of total *Dloop*, *ND4* and *16s* in wild-type and *Lrrk2^G2019S^* BMDMs with and without 4 days of 10µM ddC treatment. **M.** PI staining over a time course of AIM2 activation in wild-type and *Lrrk2^G2019S^* BMDMs +/-ddC treatment as described in L. **N.** Quantification of PI incorporation from M. at each hour. Data are expressed as a mean of three or more biological replicates with error bars depicting SEM. Statistical significance was determined using a two-tailed Student’s T test: * = p<0.05, ** = p<0.01, *** = p<0.001.

Another important mito-DAMP with connections to both inflammasome activation and cell death is mitochondrial reactive oxygen species (mito-ROS). To define the repertoire of free radicals in *Lrrk2^G2019S^* during AIM2 inflammasome activation, we used the fluorogenic probe cellROX, which emits bright green fluorescence upon oxidation and DNA binding. At 2h post-LPS/poly dA:dT treatment, we measured higher cellROX signal in *Lrrk2^G2019S^* macrophages compared to wild-type (Fig. 3D). To pinpoint the origin of this oxidative potential, we treated cells with the mitochondrial-targeted superoxide indicator mitoSOX and observed significantly higher levels of mitochondrial superoxide in *Lrrk2^G2019S^* BMDMs at 2h post-AIM2 activation (Fig. 3E). Because the transcription factor NRF2 is critical for restraining superoxide production^98^, we measured expression of NRF2 genes by RT-qPCR and found no significant differences between *Lrrk2^G2019S^* and wild-type macrophages (Fig. S3B). In the absence of transcriptional reprogramming, we hypothesized that additional mitochondrial defects could be responsible for increased mitochondrial ROS production in *Lrrk2^G2019S^* BMDMs.

### *Lrrk2^G2019S^* macrophages remain reliant on oxidative phosphorylation following engagement of pattern recognition receptors and inflammasome activation

To test whether ROS accumulation in *Lrrk2^G2019S^* BMDMs is caused by altered mitochondrial respiration, we used the Agilent Seahorse Metabolic Analyzer to measure energy production in live cells. In this assay, oxidative phosphorylation and glycolysis are assayed by oxygen consumption rate (OCR) and extracellular acidification rate (ECAR), respectively. We began by comparing mitochondrial metabolic capacity in wild-type and *Lrrk2^G2019S^* BMDMs at rest and observed no significant differences (Fig. 3F, top; Fig. 3G). However, upon LPS treatment, we observed significantly elevated spare capacity and maximal respiration in *Lrrk2^G2019S^* BMDMs compared to controls (Fig. 3F, middle, and Fig. 3G-H). This suggests that *Lrrk2^G2019S^* BMDMs fail to undergo the well-characterized metabolic shift from OXPHOS to glycolysis that results in lower OCR^99^, and instead continue to rely on OXPHOS following macrophage activation. We observed a similar phenotype—whereby the “acute response” to the addition of glucose was dampened in *Lrrk2^G2019S^* BMDMs—upon AIM2 inflammasome activation (Fig. 3F, bottom and 3I). Lastly, we noted that *Lrrk2^G2019S^* BMDMs carry out higher levels of non-mitochondrial oxygen consumption (Fig. 3J), which is generally associated with high cellular oxidase activity and ROS accumulation^100,101^. Notably, OCR defects were evident only in the initial macrophage respiratory burst, as wild-type and *Lrrk2^G2019S^* BMDMs treated with LPS for 24h undergo the OXPHOS to glycolysis transition expectedly (Fig. S3C-D). No significant phenotypic differences between wild-type and *Lrrk2^G2019S^* BMDMs were observed at the level of ECAR in response to any stimuli, suggesting that *Lrrk2^G2019S^* does not impact the glycolytic capacity of macrophages (Fig. S3C, E). Together, these data demonstrate that *Lrrk2^G2019S^* BMDMs are defective in metabolic reprogramming in response to immune stimuli. We propose this failure to shift from OXPHOS to glycolysis in *Lrrk2^G2019S^* BMDMs places additional pressure on electron transport chains, which contributes to elevated ROS levels (Fig. 3D-E).

Having observed altered mitochondrial dynamics in *D. melanogaster* expressing hLRRK2-G2019S (Fig. 1J), we asked whether mitochondria similarly failed to respond to immune stimuli in these flies. Flies were orally infected with the bacterial entomopathogen strain *Pseudomonas entomophila* (P.e.) at OD600 = 50-60 (or sucrose alone as a control). After 16-20h of infection, the fat body/adipose tissue was dissected, fixed, and immunostained to visualize mitochondria and lipid droplets as in Fig. 1J. We observed a dramatic increase in mitochondrial density upon *P.e.* infection of wild-type *D.m.* (Fig. 3K, left, mock vs. *P.e*.), which is demonstrative of the increased demand for mitochondrial output upon infection of flies, as these insects rely on a battery of energetically costly responses to eliminate enteric pathogens, including defecation^84^. In contrast, we observed no change in the density of the mitochondrial network in hLRRK2-G2019S-expressing flies in response to *P.e.* infection. Likewise, although lipid droplet size decreased in *P.e.*-infected control flies, droplet size was unchanged in hLRRK2-G2019S-expressing flies (Fig. 3K, right, mock vs. *P.e.*). These data suggest that the impact of the *Lrrk2^G2019S^* allele on the ability of mitochondria to adapt to changing energy needs during innate immune activation is evolutionarily conserved.

We next sought to link mitochondrial defects and mito-DAMP release with *Lrrk2^G2019S^* cell death. To this end, we treated cells with the nucleoside analog 2′,3′-dideoxycytidine (ddC) to inhibit mtDNA replication. Cells treated with ddC for 5 days showed a robust decrease in mtDNA copy number (*Dloop*, *ND4*, *16s*) measured by qPCR (Fig 3L). While mtDNA depletion was sufficient to rescue inflammasome-mediated cell death in *Lrrk2^G2019S^* macrophages (Fig. 3M-N; grey vs. red), it did not significantly impact PI incorporation in wild-type BMDMs (Fig. 3M-N, black vs. blue). This argues that mitochondrial metabolism and/or mtDNA release uniquely contributes to inflammasome-triggered cell death in the presence of the *Lrrk2^G2019S^* allele.

### Cell death, mitochondrial depolarization, and mitochondrial ROS accumulation in *Lrrk2^G2019S^* macrophages is GSDMD-dependent

We next sought to elucidate the molecular mechanisms responsible for uncoupling caspase-1-dependent inflammasome-triggered cell death and IL-1β release in *Lrrk2^G2019S^* macrophages. In addition to promoting cleavage and maturation of pro-IL-1β and pro-IL-18 upon inflammasome activation, caspase-1 also cleaves gasdermin-D (GSDMD) to form N-GSDMD^102,103^. N-GSDMD oligomerizes to form pores that actively translocate IL-1β and IL-18 out of the cell and cause pyroptosis^104–106^. To determine whether GSDMD contributes to *Lrrk2^G2019S^*- dependent cell death in macrophages, we measured PI incorporation after AIM2 stimulation in the presence of two GSDMD inhibitors: disulfram and necrosulfamide, which inhibits mouse GSDMD (used here), and human MLKL^107^. In the presence of both disulfram (Fig. 4A) and necrosulfamide (Fig. S4A), PI incorporation in *Lrrk2^G2019S^* macrophages returned to wild-type levels. We also observed rescue of *Lrrk2^G2019S^*-dependent cell death during inflammasome activation when we knocked down GSDMD expression using siRNAs (Fig. 4B and S4B). Together, these data pinpoint GSDMD as a critical factor in *Lrrk2^G2019S^*-dependent cell death following inflammasome activation.

**Figure 4.**
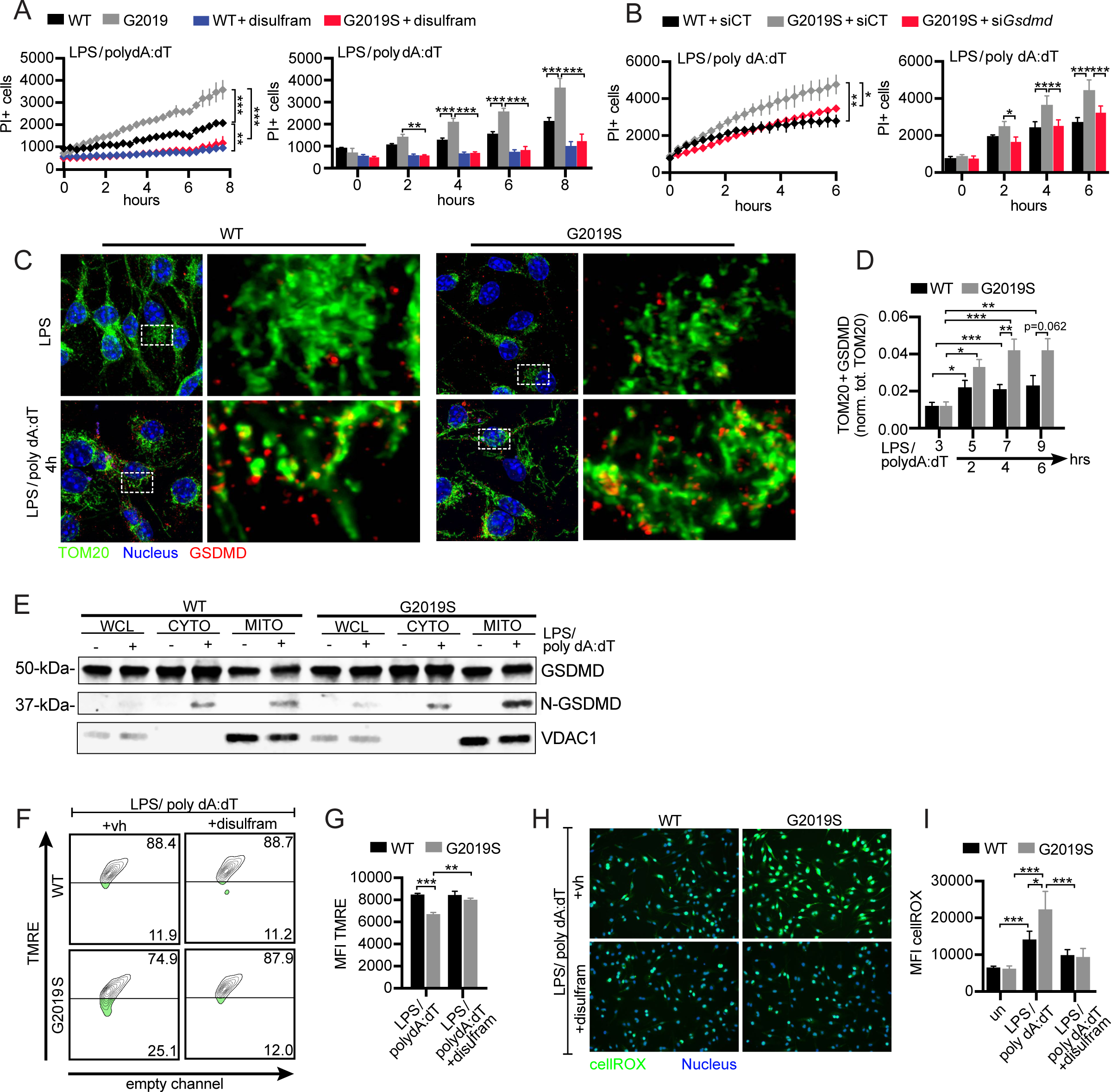
GSDMD mediates mitochondrial dysfunction and cell death during inflammasome activation in *Lrrk2^G2019S^* BMDMs. **A.** PI staining over a time course of AIM2 activation in wild-type and *Lrrk2^G2019S^* BMDMs +/-1µM disulfram at the time of priming. (right) Quantification of PI incorporation from A. at each hour **B.** PI staining over a time course of AIM2 activation in wild-type and *Lrrk2^G2019S^* BMDMs following introduction of an siRNA against *Gsdmd* or an untargeted control siRNA. (right) Quantification of PI incorporation from B. at each hour **C.** Immunofluorescence microscopy images visualizing mitochondria (anti-TOM20, green) and GSDMD (anti-GSDMD, red) of wild-type and *Lrrk2^G2019S^* BMDMs 3h post-LPS treatment (top) or 4h post-AIM2 activation. Nuclei visualized by DAPI (blue). **D.** Quantitation of TOM20+ GSDMD aggregates normalized to total TOM20 over a time course of AIM2 stimulation **E.** Association of N-GSDMD with mitochondrial network in wild-type and *Lrrk2^G2019S^* BMDMs via biochemical fractionation and immunoblot analysis. VDAC1 used as a control for mitochondrial membrane enrichment. **F.** TMRE staining of wild-type and *Lrrk2^G2019S^* BMDMs treated with 1 µM disulfram, followed by AIM2 activation for 2h. **G.** Quantitation of TMRE MFI from G. **H.** Immunofluorescence microscopy images of cellROX (green) staining in wild-type and *Lrrk2^G2019S^* BMDMs +/-1 µM disulfram 2h post-AIM2 stimulation (live nuclei staining with NucBlue). **I.** Quantitation of cellROX MFI from E. Data are expressed as a mean of three or more biological replicates with error bars depicting SEM. Statistical significance was determined using a two-tailed Student’s T test: * = p<0.05, ** = p<0.01, *** = p<0.001.

A simple explanation for GSDMD driving *Lrrk2^G2019S^* -dependent cell death is that N-GSDMD forms more pores in the plasma membrane of *Lrrk2^G2019S^* BMDMs, leading to more pyroptosis. However, we would expect that enhanced GSDMD pore formation would be concomitant with more IL-1β release, which we did not observe (Fig. 2N). Therefore, we hypothesized that GSDMD contributes to *Lrrk2^G2019S^*-dependent cell death via an alternative mechanism. Recent studies have observed that gasdermins can traffic to the mitochondria during NLRP3 inflammasome activation, which results in enhanced inflammasome activity via GSDMD-mediated mitochondrial stress and DAMP release^108^. To test whether GSDMD interaction with the mitochondrial network is altered in *Lrrk2^G2019S^* BMDMs, we used immunofluorescence microscopy with antibodies for GSDMD (red) and the mitochondria (TOM20; green) to assess GSDMD colocalization with mitochondria after AIM2 activation. Inflammasome activation, compared to LPS alone, triggered a significant accumulation of GSDMD at the mitochondria in both wild-type and *Lrrk2^G2019S^* BMDMs (Fig. 4C). Compared to wild-type, *Lrrk2^G2019S^* BMDMs had higher levels of GSDMD association at later time points following AIM2 activation (Fig. 4D). We also measured increased N-GSDMD association with mitochondrial membranes in *Lrrk2^G2019S^* BMDMs following AIM2-stimulation by immunoblot, looking for enrichment of N-GSDMD in mitochondrial membrane fractions (VDAC1-enriched) compared to cytosolic fractions (Fig. 4E). These data demonstrate that AIM2 stimulation promotes N-GSDMD mitochondrial association and that this phenomenon occurs preferentially in *Lrrk2^G2019S^* BMDMs.

Having linked lower mitochondrial membrane potential and excessive mito-ROS production to cell death in *Lrrk2^G2019S^* BMDMs (Figs. 3A and 3D), we asked whether GSDMD contributed to these defects by treating cells with the GSDMD inhibitor disulfram during AIM2 activation. Remarkably, disulfram treatment was sufficient to return both *Lrrk2^G2019S^* BMDM mitochondrial membrane potential, as measured by TMRE (Figs. 4F-G), as well as cellular ROS, as measured by cellROX (Figs. 4H-I), to wild-type levels. Together this suggests that GSDMD contributes to mitochondrial network damage and higher ROS levels in *Lrrk2^G2019S^* BMDMs.

### N-GSDMD forms pores in the macrophage mitochondrial network during AIM2 inflammasome activation

Because we observed GSDMD association with hyper-fragmented mitochondria in wild-type cells following AIM2 activation (Fig. 5A), we set out to determine whether association of N-GSDMD with mitochondrial membranes is a general phenomenon during inflammasome activation. First, we measured GSDMD association with mitochondria using flow cytometry, as in Fig. 1C-F, but this time co-staining with MitoTracker and anti-GSDMD antibodies. Consistent with our cellular fractionation experiments, which detect GSMDM on mitochondria in both wild-type and *Lrrk2^G2019S^* cells, we observed an increase in GSDMD association with mitochondria following AIM2 stimulation of wild-type BMDMs (Fig. 5B). This was dependent on GSDMD’s pore forming ability since treatment with disulfram abolished the GSDMD-mitochondrial interaction (Fig. 5B, right plot).

**Figure 5.**
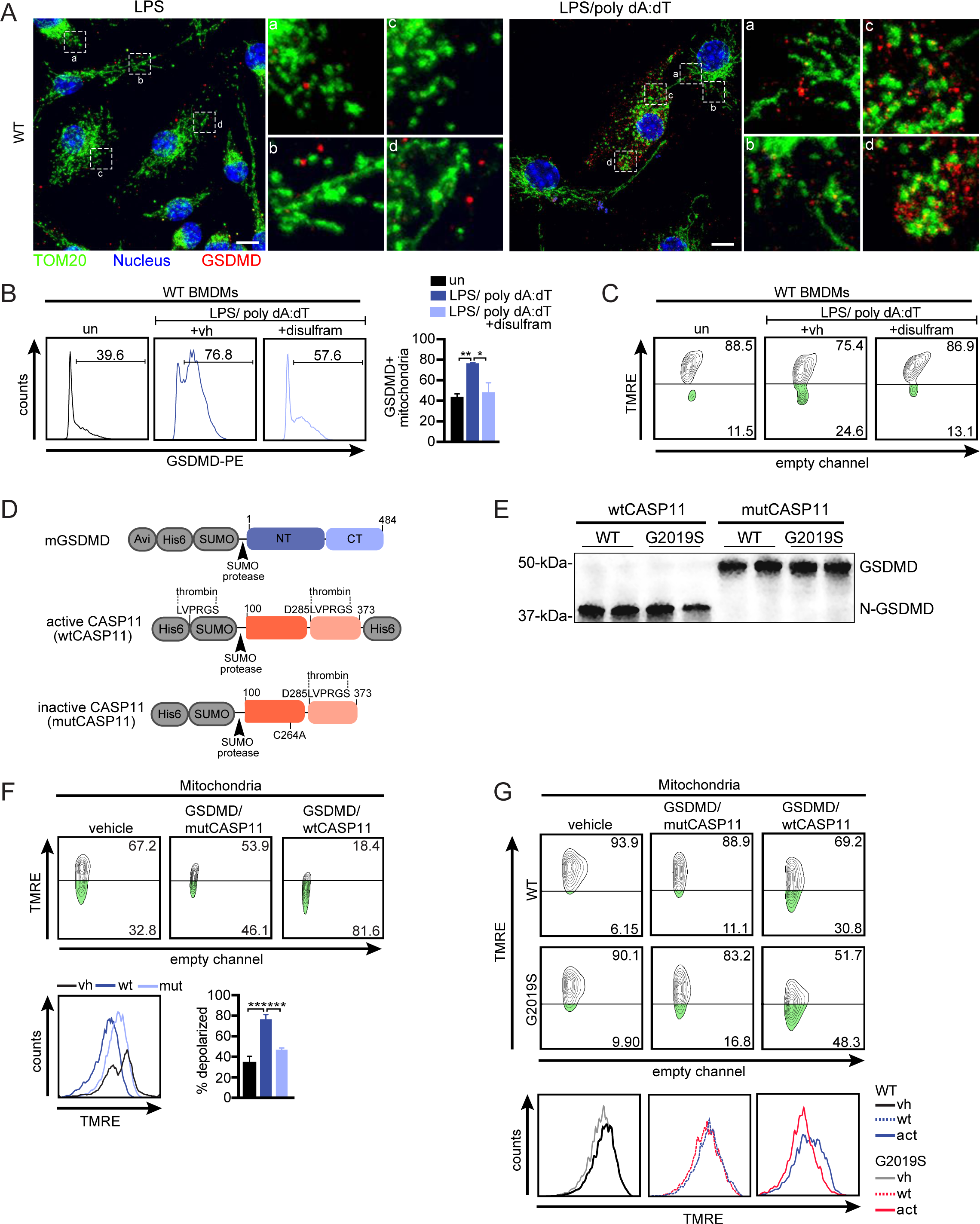
N-GSDMD directly mediates depolarization of macrophage mitochondrial membranes following AIM2 activation. **A.** Immunofluorescence microscopy images of mitochondrial network fragments in wild-type BMDMs (anti-GSDMD, red; anti-TOM20, green, nuclei visualized with DAPI). **B.** GSDMD association (anti-GSDMD PE) with the mitochondrial network as measured by flow cytometry on isolated mitochondria 4h after AIM2 inflammasome activation +/- disulfram treatment. (1 µM). **C.** TMRE staining of wild-type BMDMs treated with 1 µM disulfram, followed by AIM2 activation for 4h. **D.** Schematic representation of GSDMD, mutCASP11, and wtCASP11 recombinant proteins used in *in vitro* experiments. **E.** Immunoblot of GSDMD cleavage by recombinant CASP11. Samples run in duplicate. **F.** TMRE staining of mitochondria isolated from WT BMDMs in the presence of full length GSDMD and either the catalytically active domain (wtCASP11) or catalytic dead domain (mutCASP11) of CASP11. **G.** The same as in F. but mitochondria were isolated from wild-type and *Lrrk2^G2019S^* BMDMs. Data are expressed as a mean of three or more biological replicates with error bars depicting SEM. Statistical significance was determined using a two-tailed Student’s T test: * = p<0.05, ** = p<0.01, *** = p<0.001.

We next asked whether GSDMD association with mitochondria alters mitochondrial membrane potential in wild-type cells by measuring TMRE fluorescence over a time course. Although we did not detect significant GSDMD-dependent changes in the membrane potential of wild-type mitochondria at 2h (as we did in *Lrrk2^G2019S^* cells (Fig. 4H-1)), we observed changes in wild-type membrane potential at later time points (4h) (Fig. 5C), suggesting that GSDMD depolarizes the mitochondrial membrane in wild-type macrophages, but with slower kinetics.

To directly address the capacity of N-GSDMD to damage mitochondria, we turned to a minimal *in vitro* system. To generate N-GSDMD we combined recombinantly-expressed full length GSDMD with the catalytically active mouse CASP11 (residues 100-373, wtCASP11) or a catalytically dead form of the same protein (C254A; mutCASP11) (Fig 5D, S5A). Only wtCASP11 was capable of cleaving GSDMD (Figs. S5A,B and 5E). We then incubated mitochondria isolated from wild-type BMDMs with GSDMD/mutCASP11 or GSDMD/wtCASP11 and measured mitochondrial membrane potential via TMRE fluorescence. After 1h we observed 80% depolarization of wild-type mitochondria in the presence of GSDMD/wtCASP11 (Fig. 5F). We then sought to compare *Lrrk2^G2019S^* depolarization under these conditions. We measured no reproducible differences in N-GSDMD formation in wild-type or *Lrrk2^G2019S^* samples (Fig. 5E). However, even in this minimal system, we observe significantly more depolarization of *Lrrk2^G2019S^* in the presence of GSDMD/wtCASP11 but not GSDMD/mutCASP11 (50% depolarization after only 30min compared to 30% depolarization in the WT) (Fig. 5G). Collectively, these data demonstrate that N-GSDMD is necessary and sufficient to depolarize mitochondrial membranes upon inflammasome activation and suggest that intrinsic differences in the structure and/or composition of the mitochondrial outer membrane of cells with the *Lrrk2^G2019S^* allele render them more susceptible to this phenomenon.

### Mito-DAMP release drives necroptosis in *Lrrk2^G2019S^* macrophages

Together, our data suggest that *Lrrk2^G2019S^* macrophages die via activation of a non-pyroptotic cell death pathway that is driven by mito-DAMP release following N-GSDMD pore formation in the mitochondrial network. We next set out to test the involvement of the cell death pathways that are intimately linked to mitochondrial damage: PANoptosis, and necroptosis (Fig. 6A). PANoptosis is a newly described type of cell death in which apoptosis, pyroptosis, and necroptosis pathways are all engaged in parallel following complex inflammasome triggers like HSV or *Francisella novicida* infection^109–111^. PANoptosis is driven by formation of the PANoptosome, a complex that coordinates the efforts of AIM2, ZBP1, and pyrin to promote caspase-1-, caspase-3,7,8- and MLKL-dependent inflammatory cell death^109^. Necroptosis is another form of inflammatory programmed cell death in which the RHIM domain containing proteins RIPK1/RIPK3 activate the pore-forming protein MLKL downstream of a variety of stimuli including mtDNA release^112^, ROS and cytokine signaling^113,114^, PAMPs, and caspase inhibition.

**Figure 6.**
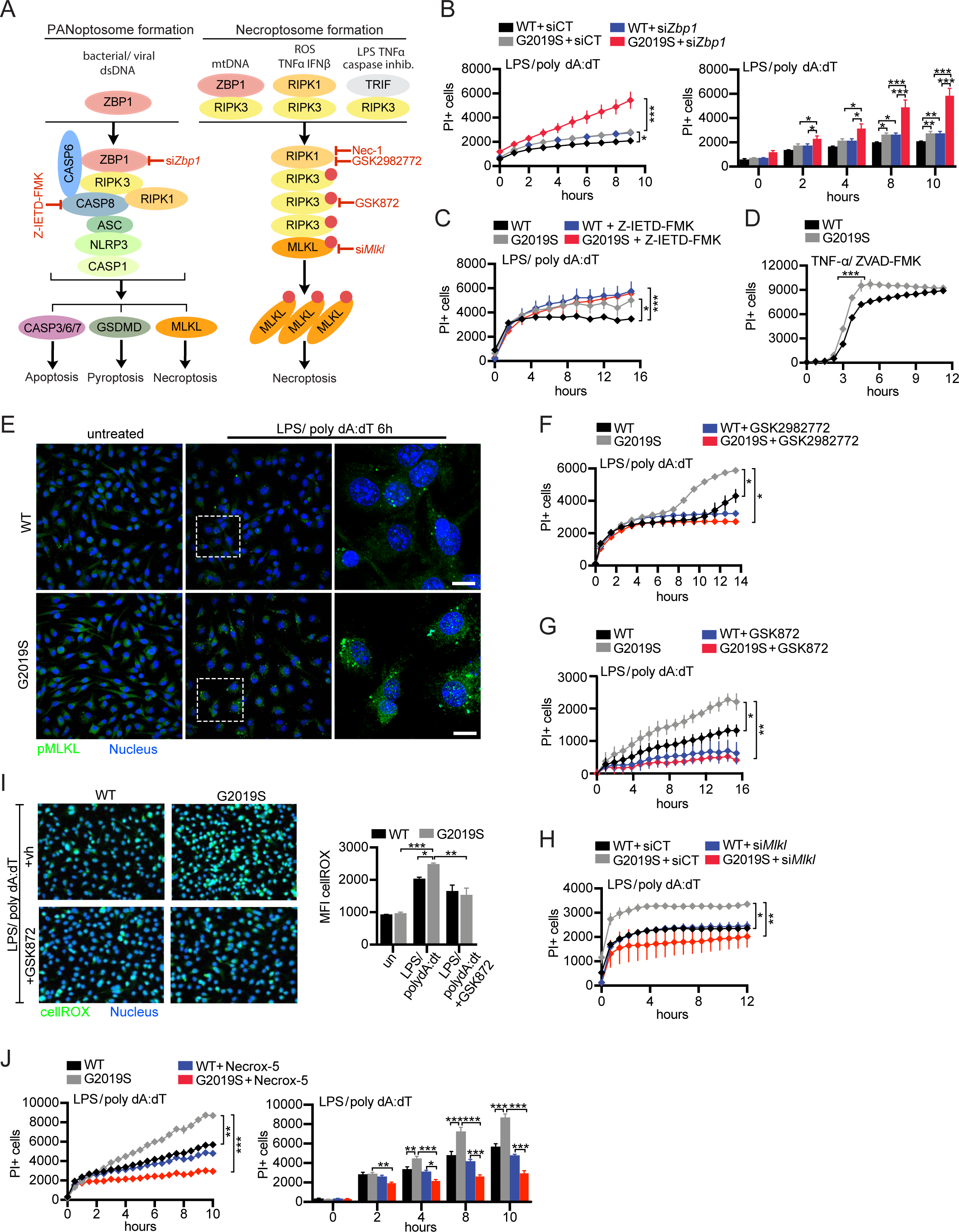
GSDMD-dependent alteration of mitochondrial homeostasis triggers RIPK1/RIPK3/MLKL-dependent necroptotic cell death in *Lrrk2^G2019S^* BMDMs. **A.** Schematic representation of PANoptosis and necroptosis signaling components. Red words/lines indicate steps in the pathway that we tested for their involvement in *Lrrk2^G2019S^* cell death **B.** PI staining over a time course of AIM2 activation in wild-type and *Lrrk2^G2019S^* BMDMs following introduction of an siRNA against *Zbp1* or an untargeted control siRNA. Quantification of PI incorporation at each hour on the right. **C.** PI staining over a time course of AIM2 activation in wild-type and *Lrrk2^G2019S^* BMDMs +/-Z-IETD-FMK. **D.** PI staining over a time course of necroptosis activation via 100 ng/ml TNF-*α*/ 50 μM ZVAD-FMK treatment in wild-type and *Lrrk2^G2019S^* BMDMs. **E.** Immunofluorescence microscopy images of pMLKL aggregation at 6h post-AIM2 stimulation in wild-type and *Lrrk2^G2019S^* BMDMs. **F.** PI staining over a time course of AIM2 activation in wild-type and *Lrrk2^G2019S^* BMDMs +/- the RIPK1 inhibitor GSK2982772. **G.** As in F but with the RIPK3 inhibitor GSK872. **H.** As in B but following introduction of an siRNA against *Mlkl*. **I.** Immunofluorescence microscopy images of wild-type and *Lrrk2^G2019S^* BMDMs stained with fluorogenic oxidative stress probe cellROX (green) and live cell nuclear stain NucBlue (blue) 2h post-AIM2 activation +/- RIPK3 inhibitor GSK872. (right) Quantification of cellROX MFI in wild-type and *Lrrk2^G2019S^* BMDMs **J**. PI staining over a time course of AIM2 activation in wild-type and *Lrrk2^G2019S^* BMDMs +/- the mitochondrial ROS scavenger Necrox-5 (25 μM). Data are expressed as a mean of three or more biological replicates with error bars depicting SEM. Statistical significance was determined using a two-tailed Student’s T test: * = p<0.05, ** = p<0.01, *** = p<0.001.

Having determined that AIM2 activation is sufficient to trigger enhanced cell death in *Lrrk2^G2019S^* macrophages, we asked whether this cell death shares characteristics of PANoptosis. To determine if ZBP1 contributes to *Lrrk2^G2019S^* -dependent enhanced cell death, we transfected wild-type and *Lrrk2^G2019S^* with siRNAs to knockdown *Zbp1* (Fig. S6A) and stimulated inflammasome activation via LPS polydA:dT. Surprisingly, we found that loss of *Zbp1* was not sufficient to rescue *Lrrk2^G2019S^*-dependent enhanced cell death; in fact, it promoted higher levels of PI incorporation in *Lrrk2^G2019S^* macrophages (Fig. 6B). We next tested the involvement of caspase-8 in *Lrrk2^G2019S^*-dependent enhanced cell death using the inhibitor Z-IETD-FMK. Despite its previous characterization as a molecular switch between different modes of cell death^115^, we found that caspase-8 had no effect on PI incorporation in *Lrrk2^G2019S^* BMDMs compared to wild-type (Fig. 6C). Because the PANoptosome involves the coordinated efforts of AIM2, ZBP1, and caspase-8^109^, these data argue that PANoptosis is not active in *Lrrk2^G2019S^* macrophages following inflammasome activation.

We next hypothesized that inflammasome activation and GSDMD pore formation drives necroptosis in *Lrrk2^G2019S^* macrophages. To implicate components of the necroptosome in *Lrrk2^G2019S^* cell death, we first asked if *Lrrk2^G2019S^* macrophages were prone to elevated death when necroptosis is directly activated. To achieve this, we treated WT and *Lrrk2^G2019S^* BMDMs with the necroptotic stimuli TNF-*α* and the pan-caspase inhibitor ZVAD-FMK (Fig. 6A, right). Consistent with a system primed for necroptotic cell death, we saw a significant shift in the rate of cell death in *Lrrk2^G2019S^* BMDMs compared to wild-type (Fig. 6D). Next, to directly test whether the end-point of necroptosis (phosphorylation and aggregation of MLKL) occurs downstream of inflammasome activation in *Lrrk2^G2019S^* macrophages, we visualized pMLKL in wild-type and *Lrrk2^G2019S^* BMDMs using immunofluorescence microscopy at 6h post-LPS/poly dA:dT treatment. Aggregates of pMLKL were readily visible in *Lrrk2^G2019S^* cells following AIM2 stimulation (Fig. 6E). To further implicate specific necroptosome components in *Lrrk2^G2019S^*-mediated cell death, we measured PI incorporation after inflammasome activation in wild-type and *Lrrk2^G2019S^* BMDMs treated with the RIPK1 inhibitors GSK2982772 and necrostatin-1 (Nec-1), treated with the RIPK3 inhibitor GSK872, or transfected with *Mlkl* siRNA (Fig. S6B). Inhibition of RIPK1 (Figs. 6F and S6C), RIPK3 (Fig. 6G), or loss of *Mlkl* expression (Fig. 6H), were each sufficient to rescue cell death in *Lrrk2^G2019S^* macrophages. Together, these results indicate that RIPK1/RIPK3/MLKL-dependent necroptosis occurs downstream of inflammasome activation in *Lrrk2^G2019S^* macrophages.

We then sought to directly implicate mitochondrial damage and mito-ROS accumulation in this pyroptosis-to-necroptosis shift. To this end, we measured cellROX after inhibiting RIPK3, and found that it was sufficient to return *Lrrk2^G2019S^* ROS levels to those of wild-type cells (Fig. 6I). This finding is consistent with previous studies showing that activated RIPK3 promotes aerobic respiration leading to ROS production through pyruvate dehydrogenase activation^116^. Treatment of cells with the mitochondrial ROS scavenger Necrox-5 also decreased *Lrrk2^G2019S^*-mediated cell death (Fig. 6J). Together, these data argue that ROS is both a consequence and a driver of necroptotic cell death in *Lrrk2^G2019S^* macrophages and provide evidence for a ROS-mediated feed-forward loop that enhances cell death and mitochondrial damage.

### Bacterial infection promotes hyperinflammation and pathogenesis in hLRRK2 G2019S flies

Having measured a dramatic defect in the ability of hLRRK2 G2019S-expressing *D. melanogaster* to upregulate mitochondrial dynamics during infection with *P. entomophila* (Fig. 3K), we next asked whether flies exhibit increased susceptibility to pathogen challenge. Briefly, flies were orally infected with *P. entomophila* via overnight feeding (20h) and survival was monitored for 10-11 days. To determine whether survival was specifically impacted by expression of the hLRRK2-G2019S allele, we infected flies expressing wild-type hLRRK2, hLRRK2 G2019S, and a kinase dead version of hLRRK2 (K1906M), alongside wild-type *w118* controls, and followed flies over the course of 10 days (Fig. 7A). Remarkably, flies ubiquitously expressing hLRRK2-G2019S were more likely to succumb to *P.e.* infection than were flies expressing hLRRK2 or the K1906M mutant allele (Fig. 7B, left). We observed a similar phenomenon when hLRRK2 expression was limited to the fat body/adipose (CGGal4), although interestingly, in that tissue, expression of hLRRK2-wild-type was sufficient to decrease survival, suggesting LRRK2 expression influences pathogenesis in a dose dependent fashion (with G2019S being a constitutively active gain-of-function allele) (Fig. 7B, right). Surprisingly, flies expressing hLRRK2-K1906M in the fat body showed enhanced survival compared to wild-type flies, suggesting that kinase dead LRRK2 promotes innate immune protection in *D. m.* (Fig. 7B, red). In *P.e*.-infected flies, levels of innate immune response genes (*Drs, Atta, and Dipt,* markers of inflammation) were higher in the presence of hLRRK2 G2019S (Fig. 7C and S7A), and hyper-inflammation has been linked with susceptibility to *P.e.* infection in flies^84^. Consistent with their increased survival, expression of the kinase dead allele (hLRRK2 G2019S-K1906M) decreased inflammation to levels below those of wild-type flies (Fig. 7C and S7A). Collectively, these data show that the *Lrrk2^G2019S^* allele promotes hyper-inflammation and hyper-susceptibility to bacterial pathogen infection in flies.

**Figure 7.**
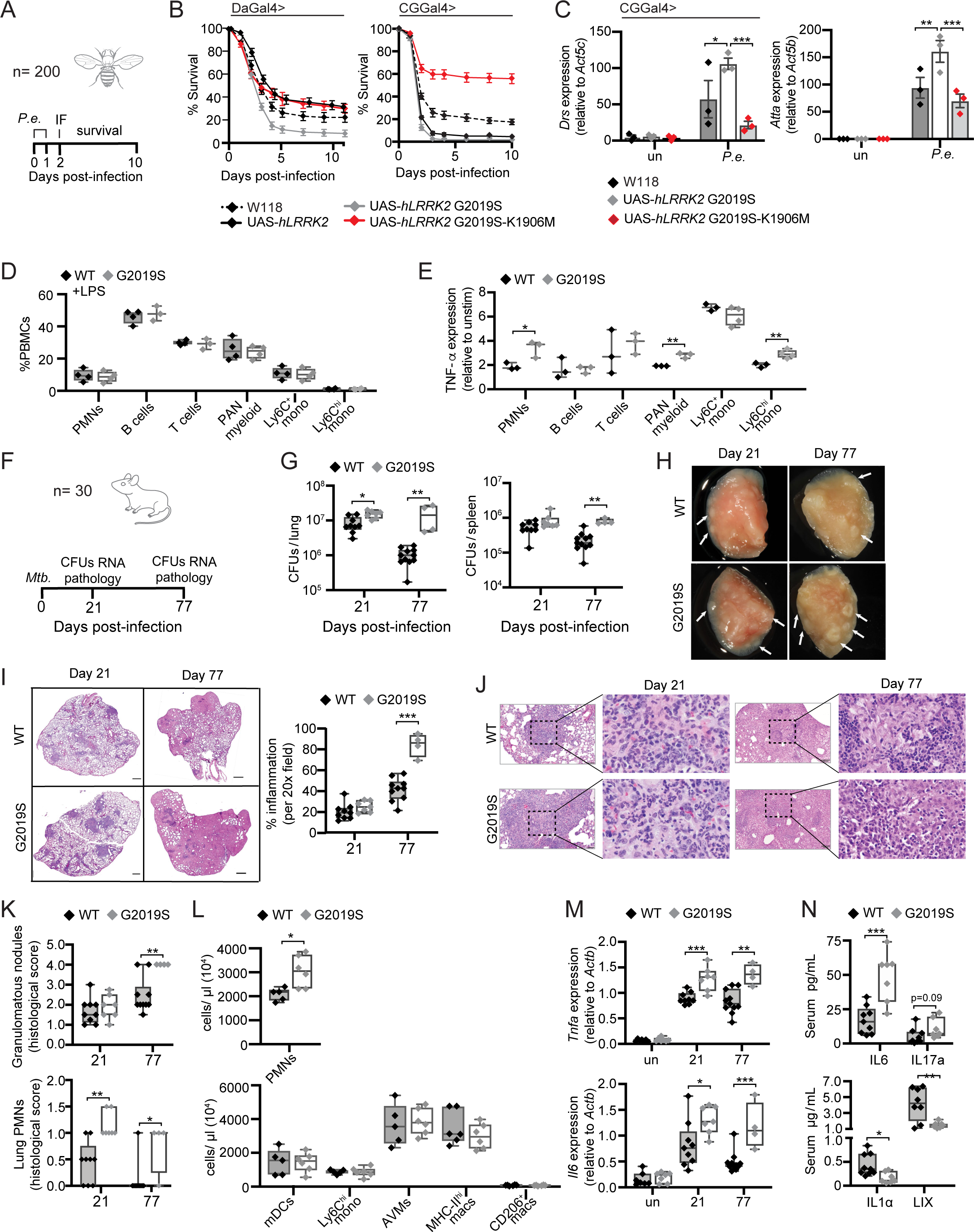
*Lrrk2^G2019S^* plays an evolutionarily conserved role in promoting hyperinflammation and susceptibility to bacterial pathogens. **A.** Schematic representation of *P.e.* infection timeline. **B.** Survival curves of wild-type (W118), hLRRK2-, hLRRK2-G2019S-, and hLRRK2 G2019S-K1906M-expressing flies (both ubiquitous expression (DaGal4>) and fat body tissue-specific expression (CGGal4>)) over a 10-day period after *P.e.* infection (20 hours). **C.** RT-qPCR of innate immune response genes (*Drs* and *Atta*) in wild-type (W1118), hLRRK2-, hLRRK2-G2019S-, and hLRRK2 G2019S-K1906M-expressing flies after *P.e.* infection (20 hours). Gene expression is shown relative to an *Act5c* control. **D.** PBMC cell population percentages in wild-type and *Lrrk2^G2019S^* mice 4 hours after stimulation with 1 µg/ml LPS as measured by flow cytometry. **E.** TNF-α protein expression in PBMCs treated as in A. TNF-α expression is shown as fold change comparing MFI in resting vs. stimulated cells. **F.** Schematic representation of Mtb infection timeline. **G.** Mtb colony forming units (CFUs) recovered from the lung and spleen of wild-type and *Lrrk2^G2019S^* infected mice at day 21 and 77 post-infection. **H.** Inflammatory nodules in the lungs of wild-type and *Lrrk2^G2019S^* Mtb-infected mice at day 21 and 77 post-infection. **I**. Hematoxylin and eosin (H&E) stain of inflammatory nodules in the lungs of wild-type and *Lrrk2^G2019S^* Mtb-infected mice 21 and 77 days after infection with Mtb. Quantification of percent inflammation on right. **J.** H&E stain of neutrophils within an inflammatory nodule in the lung of wild-type and *Lrrk2^G2019S^* Mtb-infected mice at day 21 and 77 post-infection. **K.** (top) Semiquantitative score of pulmonary inflammation with a score of 0, 1, 2, 3 or 4 based on granulomatous nodules in none, up to 25%, 26–50%, 51–75% or 76–100% of fields, respectively. (bottom) Quantification by pathology scoring of polymorphonuclear neutrophils (PMNs) in the lungs of wild-type and *Lrrk2^G2019S^* Mtb-infected mice at day 21 and 77 post-infection. **L**. Lung innate immune cell populations as measured by flow cytometry on day 21 post infection. **M**. RT-qPCR of inflammatory cytokines (*Tnfa* and *Il6*) from total RNA recovered from lung homogenates from uninfected and Mtb-infected wild-type and *Lrrk2^G2019S^* mice at day 21 and 77 post-infection. **N.** Serum cytokines IL-6, IL-17a, IL-1α, and LIX, in wild-type and *Lrrk2^G2019S^* mice measured by cytokine array (Eve Technologies) at day 21 after Mtb infection. Data are expressed as a mean of three or more biological replicates with error bars depicting SEM. Statistical significance was determined using a two-tailed Student’s T test: * = p<0.05, ** = p<0.01, *** = p<0.001.

### *Lrrk2^G2019S^* mice do not display altered immune cell populations or significantly altered ability to respond to LPS challenge *ex vivo*

To further translate our findings to models of human disease, we next asked whether mice harboring the *Lrrk2^G2019S^* allele display any differences in circulating immune cell populations. Previous characterization of *Lrrk2^G2019S^* hemizygous mice found that they display no gross motor or neuronal defects^117^ and no baseline differences in blood chemistry or white blood cell counts^118^. To elaborate on this earlier analysis, we measured populations of immune cells in blood isolated from wild-type and *Lrrk2^G2019S^* mice by flow cytometry. We found no significant differences between populations of any white blood cell in wild-type and *Lrrk2^G2019S^* mice (Figs. 7D and S7B). To determine whether isolated white blood cells differed in their ability to respond to innate immune challenge, we treated isolated blood cells with 1 µg/ml of LPS and measured expression of intracellular TNF-*α*. We found that polymorphonuclear leukocytes (PMNs), pan-myeloid lineage, and CD11b^+^ Ly6C^hi^ monocytes were inflammation prone in *Lrrk2^G2019S^* mice (Fig. 7E), when compared to unstimulated cells (Fig. S7C). This demonstrates that while certain myeloid populations are prone to hyperinflammation in response to TLR stimulation, *Lrrk2^G2019S^* mice have similar numbers of circulating immune cells compared to wild-type.

### *Lrrk2^G2019S^* promotes hyperinflammation and immunopathology during *Mycobacterium tuberculosis* infection

Because of its intracellular lifestyle, *Mycobacterium tuberculosis* has evolved an exquisite level of control over cell death pathways in macrophages. Disruption of this balance has dramatic impacts on mycobacterial disease progression in animal models, with apoptotic macrophage cell death generally being anti-bacterial and necrosis being pro-bacterial^119–126^. Given the propensity for *Lrrk2^G2019S^* macrophages to undergo programmed necrotic cell death (necroptosis) following inflammasome activation, and because GWAS correlate LRRK2 SNPs with mycobacterial susceptibility^63–65,127^, we predicted that *Lrrk2^G2019S^* mice would experience exacerbated pathogenesis in response to Mtb infection. To test this, we used our low dose aerosol infection model to administer approximately 100 Mtb bacilli per mouse to a cohort of both male and female wild-type and *Lrrk2^G2019S^* mice. We sacrificed mice at day 21, which provides a snapshot of the innate immune response to Mtb infection, and at day 77, which captures both innate and adaptive responses (Fig. 7F). We observed dramatically higher bacterial burdens in the lungs of *Lrrk2^G2019S^* mice compared to wild-type at day 77, with a moderate but significant increase seen as early as day 21 (Fig. 7G, left). Significantly higher CFUs were also found in the spleens of *Lrrk2^G2019S^* mice at day 77 (Fig. 7G, right). These data demonstrate that expression of the *Lrrk2^G2019S^* allele in mice creates a favorable niche for Mtb survival and replication.

We next assessed inflammation and cellular infiltration in the lungs of wild-type and *Lrrk2^G2019S^* mice over the course of Mtb infection. Gross examination upon tissue collection revealed more extensive inflammatory nodules within the lungs of *Lrrk2^G2019S^* mice compared to wild-type controls, on both 21 and 77 days post-infection (Fig. 7H). Histologically, lungs from *Lrrk2^G2019S^* mice trended towards greater inflammatory infiltrates at day 21 and by day 77, *Lrrk2^G2019S^* lungs were almost completely obscured by coalescing inflammatory cell infiltrates (compared to about half of wild-type lung tissue) (Fig. 7I). To gain additional insight into the nature of this dramatic lung inflammation, we analyzed the H&E staining of lung sections for immune cell infiltrates and found increased numbers of granulomatous nodules and increased PMNs (mainly neutrophils) over the course of Mtb infection in *Lrrk2^G2019S^* mice (Fig. 7J-K). This increase was particularly striking at day 77 when no wild-type mice scored positive for PMNs but several *Lrrk2^G2019S^* mice still had signs of neutrophil influx (Fig. 7K, bottom). We also observed increased neutrophil (CD45^+^ B220^-^ CD11b^+^ Ly6G^+^) recruitment to the lungs of Mtb-infected *Lrrk2^G2019S^* mice at day 21 post infection using flow cytometry (Fig. 7L, top). Notably, we did not measure differences in other innate immune cell types (e.g. alveolar macrophages, M1 macrophages (MHC-II^hi^), M2 macrophages (CD206+), and myeloid dendritic cells) in the lungs of wild-type and *Lrrk2^G2019S^* mice at this time point(Fig. 7L, bottom).

Consistent with higher neutrophil recruitment, we detected elevated expression of pro-inflammatory cytokine transcripts (*Tnfa*, *Il6*, *Il1b*, *Ifng*, and *Cxcl1*) in the lungs of *Lrrk2^G2019S^* Mtb-infected mice compared to wild-type controls at early and late infection time points (Fig. 7M and Fig. S7D). Type I IFN transcripts were also higher in the lungs of *Lrrk2^G2019S^* mice during Mtb infection (Fig S7D), paralleling what we observed in *Lrrk2^G2019S^* macrophages following AIM2 activation (Fig. S3A).

To define the nature of the circulating immune milieu in wild-type and *Lrrk2^G2019S^* Mtb-infected mice, we measured levels of key inflammatory cytokines and chemokines in serum via cytokine array. Curiously, despite dramatic differences in the lungs, the circulating levels of many soluble immune mediators were similar in wild-type and *Lrrk2^G2019S^* Mtb-infected mice, suggesting that local responses are the primary drivers of Mtb pathogenesis in these mice (Fig. S7E). We did, however, find significantly higher levels of IL-6 and IL-17a and lower levels of IL-1α and LIX (Fig. 7N). High IL-6 levels are consistent with cells favoring necroptotic over pyroptotic cell death, as are low levels of IL-1α, which can be released by monocytes along with IL-1β during pyroptosis^128^. Taken together, these studies suggest that in in *Lrrk2^G2019S^* mice, Mtb infection causes local hyperinflammatory responses in the lung, leading to immunopathology and poor infection outcomes.

## DISCUSSION

Cellular surveillance of mitochondrial integrity is a key checkpoint in the control of programmed cell death and release of inflammatory mediators. Here, we demonstrate that the common human disease-associated allele *Lrrk2^G2019S^* compromises mitochondrial function in macrophages. We report that upon infection with intracellular bacterial pathogens like Mtb, *Lrrk2^G2019S^* primary murine macrophages die in a manner consistent with RIPK1/RIPK3/MLKL-dependent necroptosis despite upstream activation of inflammasome-dependent pyroptosis. This “switch” relies in large part on mito-DAMP release mediated by permeabilization of the mitochondrial network by the pore-forming pyroptotic protein GSDMD, which directly associates with and depolarizes mitochondrial membranes. In mice and flies, *Lrrk2^G2019S^* expression limits the ability of cells and tissues to make energetic adjustments in response to immune stimuli, resulting in hyperinflammation and enhanced susceptibility to bacterial infection. These findings argue that LRRK2’s role in regulating immunity via controlling mitochondrial homeostasis is evolutionarily conserved and that in the presence of underlying mitochondrial dysfunction, infection with bacterial pathogens can elicit distinct, deleterious immune outcomes. Consequently, these studies necessitate a reworking of how we define cell death and innate immune “inputs” and “outputs” in the face of mitochondrial mutations and stress.

Our data suggest that GSDMD skews cell death modalities in *Lrrk2^G2019S^* macrophages by disrupting mitochondrial homeostasis. Previous research has shown that both N-GSDME and N-GSDMD can augment caspase 3/7-mediated apoptotic cell death in cancer cells by promoting release of proapoptotic proteins like cytochrome C from mitochondria^108^. GSDMD has also been shown to target mitochondria following intracellular LPS-dependent activation of caspase 11 in endothelial cells^129^. In this context, N-GSDMD promotes mtDNA release and cGAS/STING activation in the absence of cell death, suggesting that redistribution of GSDMD to and from the mitochondria may constitute an important node in regulating cell death outcomes. Here, we show that N-GSDMD decreases mitochondrial membrane potential and increases mito-DAMP release following AIM2 activation of macrophages. Based on these findings, we conclude that GSDMD is an unappreciated player in controlling mitochondrial homeostasis following caspase activation. Understanding why such a mechanism exists in macrophages and the extent to which it occurs in other cell types or downstream of other cell death stimuli will require further study. It is possible that, as proposed by Rogers et al., N-GSDMD forms pores in mitochondria to promote release of factors like cytochrome C to amplify cell death signals^108^. It is also possible that GSDMD association with the mitochondria is another carry over from the Rickettsial origins of mitochondria, as N-GSDMD can bind to and kill bacteria in *in vitro* cultures^104^.

We further demonstrate that GSDMD mitochondrial association and depolarization occurs more readily in *Lrrk2^G2019S^* mitochondria (Figs. 4-5). We propose that the enhanced ability of N-GSDMD to damage *Lrrk2^G2019S^* mitochondria stems directly from mitochondria stress and/or changes to the mitochondrial membrane attributed to the *Lrrk2^G2019S^* allele. It is possible that mitochondrial hyper fission confers susceptibility to GSDMD pore formation, much in the same way that BAX-dependent membrane permeabilization has mitochondrial size and shape requirements^130^. Alternatively, fragmentation of the mitochondrial network may simply increase surface area for N-GSDMD to bind to. In support of these hypotheses, our experiments visualizing GSDMD association with the mitochondrial network show marked enrichment for GSDMD around smaller TOM20+ fragments (Fig. 5A).

It is also possible that *Lrrk2^G2019S^* alters cardiolipin accessibility, which has been shown to interact with GSDMD^105^. Experiments in yeast and flies have shown that the assembly of oxidative phosphorylation complexes promotes cardiolipin remodeling and stabilization^131^; perhaps cardiolipin accumulation is a side-effect of the alterations in OXPHOS in *Lrrk2^G2019S^* macrophages. It is also possible that ROS accumulation in *Lrrk2^G2019S^* macrophages promotes oxidation of cardiolipin, which has previously been connected to increasing permeability of the mitochondrial outer membrane^132^. Alternatively, cardiolipin translocation to the outer membrane, which occurs in response to mitochondrial stress^133^ may be enhanced in *Lrrk2^G2019S^* cells. Although the precise mechanism through which *Lrrk2^G2019S^* confers mitochondrial susceptibility to GSDMD pore formation remains undetermined, it is likely that other mitochondrial mutations and stresses could enable similar phenomena.

In the presence of the *Lrrk2^G2019S^* allele, GSDMD switches from an executioner of pyroptosis to an enhancer of necroptosis downstream of inflammasome activation. This discovery lends additional support to an emerging model wherein flexibility between the use of various cell death proteins evolved as a host adaptation against intracellular pathogens that suppress or subvert one or more of these mechanisms. For example, in the absence of GSDMD, caspase-1 can mediate apoptotic cell death via the Bid-caspase-9-caspase-3 axis^134^. Likewise, caspase-8 frequently “replaces” caspase-1 in ASC complexes during bacterial infection, as well as in cases when key components of the cell death machinery are genetically ablated or pharmacologically inhibited^115,135–138^. Building on these foundational studies, our work argues that cell death proteins can stand in for each other and elicit dramatically different cell death/immune outcomes in the context of mitochondrial dysfunction stemming from a human SNP. It is tempting to speculate that reprogramming of cell death modalities is triggered by mitochondrial dysfunction due to the ancient bacterial origin of the mitochondrion, despite these mechanisms having primarily evolved as a way for cells to overcome manipulation by intracellular pathogens.

Our initial interest in LRRK2 was borne out of numerous GWAS linking LRRK2 SNPs to *Mycobacterium leprae* (Mlep) susceptibility^63–65,127^. Due to their intracellular lifestyle, both Mlep and Mtb rely heavily on certain types of cell death to ensure their survival and spread. The balance between cell death modalities has been intensely studied in the context of Mtb, where necrotic cell death is associated with macrophage escape and spread and apoptotic cell death limits Mtb pathogenesis^97,119,122,139^. Previous studies concluded that necroptotic signaling components like RIPK3 and MLKL do not significantly contribute to cell death and/or immune outcomes in Mtb-infected macrophages *ex vivo* or during *in vivo* infection of *Mlkl^-/-^* or *Ripk3^-/-^* mice^125,140,141^. Given the degree of flexibility between cell death sensors and effectors, it is likely that RIPK3/MLKL-dependent cell death may only prioritized be in certain cellular contexts, rendering RIPK3 and MLKL dispensable in “normal” Mtb infection. It will be important to revisit the role of RIPK1/RIPK3/MLKL necroptosis during Mtb infection in additional models of mitochondrial dysfunction and stress, especially in light of therapeutic potential offered by several drugs that limit lung pathology and inflammation during Mtb infection by inhibiting necrotic cell death^142^.

The major consequence of the *Lrrk2^G2019S^* allele during Mtb infection *in vivo* is massive neutrophil infiltration, likely triggered by increased release of inflammatory DAMPs following necroptosis of Mtb-infected macrophages. There is emerging evidence that RIPK1/RIPK3/MLKL-associated necroptosis drives recruitment of neutrophils *in vivo*^143–145^ and PMN infiltration is associated with increased mycobacterial loads and immunopathology in experimental models and TB patients^146,147^. Our data are consistent with a model wherein a pyroptosis-to-necroptosis shift in Mtb-infected *Lrrk2^G2019S^* macrophages promotes excessive neutrophil infiltration and hyper-inflammation in the lungs, which is associated with poor Mtb disease outcomes in *Lrrk2^G2019S^* mice. Curiously, the same type of hyperinflammation may be protective in the context of other organisms as evidenced by *Lrrk2^G2019S^* conferring protection against *S*. Typhimurium (ST) in a sepsis model^74^, possibly due to enhanced control of ST by neutrophil influx.

Another outstanding question surrounds the nature of the inflammasome trigger that drives necroptotic cell death in *Lrrk2^G2019S^* macrophages. While we show that direct AIM2 stimulation via LPS priming followed by poly dA:dT transfection is sufficient to induce cleavage a GSDMD-dependent shift from pyroptosis to necroptosis, it is likely that other inflammasomes and/or other inflammasome triggers that activate GSDMD may accomplish the same thing. In support of this is our initial observation that cell death is enhanced during infection with ST infection, which activates the NLRC4 and NLRC3 inflammasomes, but not AIM2^148^. Interestingly, LRRK2 has previously been implicated in promoting activation of the NLRC4 inflammasome during ST infection by phosphorylating NLRC4 at Ser533^149^. It is possible that GSDMD pore formation and/or mitochondrial stress are at play during NLRC4 inflammasome activation and that many immune triggers could promote cell death via a pyroptosis-to-necroptosis switch in *Lrrk2^G2019S^* macrophages.

Despite having connected a number of LRRK2-dependent mitochondrial defects to the GSDMD-dependent necroptotic cell death we describe here, the precise mechanistic contribution of LRRK2, and LRRK2 kinase activity to the death of *Lrrk2^G2019S^* macrophages, is still not entirely clear. One intriguing possibility is that the ability of *Lrrk2^G2019S^* to push cells towards RIPK3-dependent necroptotic cell death is driven in part by LRRK2 itself being a RIP (receptor interacting protein) kinase family member (LRRK2 is sometimes annotated as RIPK7)^150^. A recent siRNA screen identified LRRK2 as a positive regulator of RIPK1-dependent apoptosis in a photoreceptor cell line and in MEFs^151^ and there is evidence that LRRK2 can co-immunoprecipitate with RIPK1 and FADD in HEK293T cells^152^. Via these interactions, perhaps wild-type LRRK2 contributes to apoptosis regulation via RIPK1/FADD/CASP8, while *Lrrk2^G2019S^* alters the composition or activation of these complexes to favor RIPK1/RIPK3-mediated necroptosis. It is also possible that *Lrrk2^G2019S^* directly phosphorylates RIPK1, RIPK3, and/or MLKL via its constitutively active kinase domain to promote necroptosome formation. Across insect genomes, one can identify innate immune receptors, adaptors, and transcription factors that encode RHIM-like motifs (like mammalian RIPKs) but lack kinase domains (unlike mammalian RIPKs). This includes the innate immune receptors/sensors PGRP-LE/LC, the innate immune adaptor IMD (immune deficiency), and the NF-kB transcription factor Relish in *Drosophila melanogaster*^153^. It is conceivable that LRRK2 may act as an ancient, and evolutionarily conserved, kinase bridge (modulating phosphorylation and activity of RHIM-containing innate immune proteins) to adjust innate immune responses in insects as well as in mammals. While it is intriguing that key LRRK2-dependent phenotypes are conserved between flies and mammals (i.e. mitochondrial network disruption, failure to metabolically adapt to innate stimuli, and hyperinflammation), additional studies are needed to determine the link, if any, between LRRK2 mitochondrial disruption and immune outcomes in the fly.

Lastly, as mutations in LRRK2 are notoriously associated with both inherited and sporadic PD, it will be important to translate our findings to cells in the CNS. While disease etiology of PD is complex and poorly understood, it has long been appreciated that the major debilitating symptoms are caused by the death of motor neurons in the substantia nigra-pars compacta (SNpc), making cell death a critical part of disease progression. While apoptosis is believed to be the core mechanism controlling neuron death during PD (reviewed in ^154^), recent analysis of human postmortem brain tissue and mouse models of the disease show activation of key components of the necroptosome machinery in dying motor neurons of the SNpc^155^. Consistent with this, inhibition of RIPK1 via necrostatin-1 rescues dopaminergic neuron loss in a mouse model of PD (MPTP treatment), suggesting a role for RIPK1 in neuronal cell death that could be modulated in the presence of the *Lrrk2^G2019S^* allele^156,157^. Furthermore, AIM2 inflammasome activity and GSDMD have been shown to play an important role in surveillance and prevention of DNA damage accumulation during neuronal development^158^. These studies argue that regulation of proinflammatory cell death pathways is critical for shaping and maintaining a healthy central nervous system. We propose that reprogramming cell death modalities via dysregulation of mitochondrial homeostasis constitutes a major way in which mutations in genes like *LRRK2* trigger and/or exacerbate human disease both in the CNS and in the periphery.

## ACKNOWLEDGEMENTS

We would like to thank the members of the Patrick and Watson labs for their helpful discussions related to experiment conceptualization and interpretation and for their help in preparing this manuscript. We would like to thank C. Elliott and W. Smit for providing us with the UAS-hLRRK2-G2019S and UAS-hLRRK2-G2019S-K1906M *Drosophila* and Dr. Raquel Sitcheran and Michael Kamradt for technical guidance in developing mitochondrial flow cytometry. We thank Malea Murphy at the Integrated Microscopy and Imaging Laboratory at TAMU COM as well as Robbie Moore and the COM-CAF core facilities at TAMU COM for their technical expertise. This work was supported by funds from the Michael J. Fox Foundation for Parkinson’s Research, Grant 12185 (to R.O.W.), the National Institutes of Health, grant R01AI125512 (to R.O.W.), and the Texas A&M Clinical Science and Translational Research (CSTR) Pilot Grant Program (to R.O.W.). Additional funding was provided by the Parkinson’s Foundation Postdoctoral Fellowship (to C.G.W.), NIH training grant 5T32OD011083-10 and the Texas A&M CVM Postdoctoral Trainee Research Training Grant (to K.J.V.).

## AUTHOR CONTRIBUTIONS

Conceptualization, K.L.P., R.O.W., C.G.W.; Investigation, C.G.W., X.Z., E.M., S.L.B., K.J.V., A.K.C., J.J.V., C.J.M., B.Z., and R.O.W.; Methodology, C.G.W., E.M., S.L.B., K.J.V., X.Z., J.K., P.L., A.P.W., R.O.W., and K.L.P.; Writing, K.L.P., C.G.W., R.O.W., J.K., X.Z.; Visualization, C.G.W., R.O.W., J.K., and S.L.B.; Funding acquisition, R.O.W., K.L.P., C.G.W., and K.J.V.; Supervision, R.O.W. and K.L.P.

## MATERIALS AND METHODS

### Mouse husbandry and strains

*Lrrk2^G2019S^* mice (B6.Cg-Tg(Lrrk2*G2019S)2Yue/J, also known as FLAG-LRRK2-G2019S, stock # 012467 were purchased from Jackson Laboratories (Bar Harbor, ME). The *Lrrk2^G2019S^* strain has been maintained with filial breeding on a C57BL6/J background. All mice used in experiments were compared to age- and sex-matched controls. Mice used to generate BMDMs were between 10-16 weeks old. Mice were infected with Mtb at 10-12 weeks. Embryos used to make primary MEFs were 14.5 days post-coitum. All animals were housed, bred, and studied at Texas A&M Health Science Center under approved Institutional Care and Use Committee guidelines.

### Bacterial culture and macrophage infections

#### M. tuberculosis

The Erdman strain was used for all *M. tuberculosis* infections. Low passage lab stocks were thawed for each experiment to ensure virulence was preserved. *M. tuberculosis* was cultured in roller bottles at 37°C in Middlebrook 7H9 broth (BD Biosciences) supplemented with 10% OADC (BD Biosciences), 0.5% glycerol (Fisher), and 0.1% Tween-80 (Fisher) or on 7H10 plates. All work with *M. tuberculosis* was performed under Biosafety level 3 containment using procedures approved by the Texas A&M University Institutional Biosafety Committee. To prepare the inoculum, bacteria grown to log phase (OD 0.6-0.8) were spun at low speed (500 g) to remove clumps, and then pelleted and washed with 1X PBS twice. Resuspended bacteria were briefly sonicated and spun at low speed once again to further remove clumps. The bacteria were diluted in DMEM (Hyclone) + 10% horse serum (Gibco) *in vitro* infections, or 1X PBS *in vivo* infections.

#### M. marinum

Low passage glycerol lab stocks of *M. marinum* were grown in 7H9 broth (BD Biosciences) supplemented with 10% OADC (BD Biosciences), 0.5% glycerol (Fisher), and 0.1% Tween-80 (Fisher). Cultures were grown at 30°C. To prepare the inoculum, cultures in log phase (OD 0.6-0.8) were spun at 2700 rcf for 10 min and washed in 1X PBS. Bacteria were pulled through a syringe with a 26 gauge needle 3 times to create single cell suspension. Bacteria were then centrifuged twice for 2 min at 2000 rcf to remove clumps. The bacteria were diluted in DMEM (Hyclone) + 10% horse serum (Gibco) and added to cells. For cell death infection assays with *M. marinum* cells were performed at 32°C.

#### L. monocytogenes

Low passage glycerol lab stocks of *L. monocytogenes* strain 10304s were streaked onto BHI agar plates and incubated at 37 °C overnight. 5-10 colonies were inoculated into 2.5 ml BHI and incubated at 30°C under static conditions overnight. To prepare the inoculum, overnight cultures were diluted 1:5 in fresh BHI, grown to log phase (OD 0.5-1.0) at 30°C. Upon reaching mid-log phase, 1 mL of bacteria were pelleted at 5000 rpm for 3 minutes and washed twice with 1X PBS. Bacteria were diluted in DMEM (Hyclone).

#### *S. enterica* (ser. Typhimirium)

Salmonella enterica serovar Typhimurium (SL1344) was obtained from Helene Andrews-Polymenis, TAMHSC. S. T. stocks were streaked out on LB plates. Overnight cultures of S. Typhimurium were grown in LB broth containing 0.3M NaCl and grown at 37 °C until they reached an OD600 of 0.9. On the day of infection cultures were diluted 1:20 on day of infection. Once cultures had reached mid-log phase, 1 ml of bacteria were pelleted at 5000 rpm for 3 minutes and washed twice with 1X PBS. Bacteria were diluted in DMEM (Hyclone).

### Primary Cell Culture

Bone marrow derived macrophages (BMDMs) were differentiated from BM cells isolated by washing mouse femurs with 10 ml DMEM. Cells were then centrifuged for 5 min at 400 rcf and resuspended in BMDM media (DMEM, 20% FBS (Millipore), 1mM sodium pyruvate (Lonza), 10% MCSF conditioned media (Waston lab)). BM cells were counted and plated at 5x10^6^ in 15 cm non-TC treated dishes in 30 ml complete BMDM media. Cells were fed with an additional 15 ml of BMDM media on day 3. Cells were harvested on day 7 with 1 X PBS EDTA (Lonza).

Mouse embryonic fibroblasts (MEFs) were isolated from embryos. Briefly, embryos were dissected from yolk sacs, washed 2 times with cold 1X PBS, decapitated, and peritoneal contents were removed. Headless embryos were disagreggated in cold 0.05% trypsin-EDTA (Lonza) and incubated on ice for 20 min., followed by incubation at 37 °C for an additional 20 min. Cells were then DNAse treated with 4 ml dissagregation media (DMEM, 10% FBS, 100 μg/ml DNAse (Worthington)) for 20 min at 37 °C. Supernatants were isolated and spun at 1000 rpm for 5 min. Cells were resuspended in complete MEF media (DMEM, 10% FBS, 1mM sodium pyruvate), and plated in 15 cm TC-treated dishes 1 dish per embryo. MEFs were allowed to expand for 2-3 days before harvest with Trypsin 0.05% EDTA.

### siRNA knockdown in BMDMs

On day 5 of differentiation BMDMs were reseeded at 0.35x10^6^ cells/ well in triplicate in 12-well non-tissue culture treated plates. The following day cells were transfected using Fugene SI reagent and 10 μM of siRNA stock. Cells were incubated for 24 hours in transfection media, then allowed to rest for 48h at 37 °C prior to downstream survival experiments.

### Cell stimulation with innate immune agonists

BMDMs were plated in 96-well half area plates at 2.5x10^4^ cells/well, 12-well plates at 5x10^5^ cells/well, 6-well plates at 1x10^6^ cells/well. To analyze upregulation of ISGs and NFkB associated genes, cells were stimulated for 4h. with 1 μM CL097 (Invivogen), 100 ng/mL LPS (Invivogen), 100 ng/mL Pam3CSK4 (Invivogen). Cells were transfected for 4h. with 1 μg/mL ISD (Watson lab), 1 μg/mL poly I:C (Invivogen), 1μg/mL cGAMP (Invivogen) using lipofectamine reagent (ThermoFisher). Cells were transfected for 4h. with 1 μM CpG 2395 (IDT) using Gene Juice (EMD Millipore). To analyze AIM2 inflammasome activation cells were stimulated with 10 ng/mL LPS (Invivogen), for 3h followed by 1 μg/mL poly dA:dT (Invivogen) transfected with lipofectamine (Thermo Fisher). For rescue assays cells were treated at the time of LPS priming with 10 μM AC-YVAD-CMK (Invivogen), 1 mM GNE9605 (Cayman Chemicals), 1μM disulfram (Sigma), 20μM necrosulfamide (Calbiochem), 10 μM necrostatin-1 (Calbiochem), 10 μM GSK2982772 (Calbiochem), 1 mM GSK872 (Calbiochem), For necroptosis induction cells were treated with 100ng/ml TNF-α (Peprotech) and 50μM Z-VAD-FMK (Sigma).

### Gene expression analysis by qRT-PCR

For mammalian tissue and cells, RNA was isolated using Direct-zol RNAeasy kits (Zymogen). cDNA was synthesized with BioRad iScript Direct Synthesis kits (BioRad) per manufacturer’s protocol. qRT-PCR was performed in triplicate wells using PowerUp SYBR Green Master Mix. Data was analyzed on a QuantStudio 6 Real-Time PCR System (Applied Biosystems), and quantification of gene expression was performed using a standard curve and normalized to *Actb* expression levels. For insect tissues, RNA from intact fly carcass (containing mostly fat body) was extracted using Trizol as per manufacturer’s protocol. cDNA was synthesized using Superscript III (Invitrogen). qRT-PCR was performed using SYBR Green, the Applied Biosystems StepOnePlus Real-Time PCR systems. Results are average ± standard error of at least three independent samples, and quantification of gene expression levels calculated using the ΔCt method and normalized to *Act5C* expression levels.

### Seahorse metabolic assays

Seahorse XF mito stress test kits and cartridges were prepared per Agilent’s protocols and analyzed on either an Agilent Seahorse XF 96-well analyzer. The day before the assay BMDMs were seeded at 5 x 10^4^ cells/well overnight. For inflammasome activation on the day of the assay cells were treated with 10 ng/mL LPS (Invivogen), followed by 1 ug/mL poly dA:dT (Invivogen) 3h later. After 2h of AIM2 activation cells were processed per manufacturer’s directions and analyzed using the Agilent Seahorse Mito Stress Test kit (Agilent). For overnight stimulation with LPS cells were treated 2h after plating with 10 ng/mL LPS and incubated overnight at 37°C.

### Flow cytometry

For submicron analysis of mitochondria by flow cytometry, cells were lifted off non-tissue culture treated plates and added to a 96-well V bottom plate. Cells were pelleted by centrifugation at 400 rcf for 3 min. Cells were resuspended in 1X PBS 2% FBS 200nM MitoTracker green (Invitrogen) and allowed to stain for 15 min at 37 °C. Cells were then washed once with PBS 2% FBS and were resuspend in ice cold mitoFLOW buffer containing 300mM sucrose, 10mM Tris (pH 7.4), 0.5mM EDTA, and 1X Halt Protease Inhibitor Cocktail. Lysis was performed by vortexing cells for 3 min followed by removal or debris by centrifugation at 400 rcf for 5 min at 4 °C. For antibody labeling samples were centrifuged at 12,000 rcf for 10 min at 4 °C and resuspend in 50 uL blocking buffer (5% BSA in mitoFLOW buffer),and incubated on ice for 15 min. Blocking was followed by an additional 20 min incubation on ice with antibodies of interest (GSDMD-PE 1:500, Abcam). Mitochondria were washed 2X in mitoFLOW buffer and were analyzed on an LSR Fortessa X20 (BD Biosciences). Flow-Jo software was used for post-acquisition analysis MitoTracker green was used to gate on mitochondria.

For JC-1 assays to assess mitochondrial membrane potential, cells were lifted from culture plates with 1X PBS + EDTA (BMDMs, RAW 264.7). Single cell suspensions were made in 1X PBS 4% FBS. JC-1 dye was sonicated for 5 min with 30 second intervals. Cells were stained for 30 min at 37°C in 1 µM JC-1 dye washed twice in PBS 4% FBS and analyzed on an LSR Fortessa X20 (BD Biosciences). Flow-Jo software was used for post-acquisition analysis. Aggregates were measured under Texas Red (610/20 600LP) and monomers under FITC (525/50 505LP). To assess mitochondrial membrane potential under stress, cells were treated for 3h with 2.5 µM rotenone prior to being lifted off the culture plates. 5 µM ATP was then added for 5, or 30 min. As a positive control 50 µM FCCP was added for 15 min.

For TMRE assays to assess mitochondrial membrane potential, cells were lifted from culture plates with 1X PBS EDTA (BMDMs). Single cell suspensions were made in 1X PBS 4% FBS. Cells were stained for 20 min at 37°C in 25 nM TMRE dye, washed 1X in PBS 4% FBS and analyzed on an LSR Fortessa X20 (BD Biosciences). Flow- Jo software was used for post-acquisition analysis. Fluorescence was measured under PE (585/15). To assess mitochondrial membrane potential during inflammasome activation cells were stimulated with 10 ng/mL LPS for 3h followed by poly dA:dT for an additional 2h prior to being lifted off the plates.

For cell death and apoptosis assays cells, were stimulated with 10 ng/ml LPS for 3h followed by 1 ug/mL poly dA:dT, or 50uM Etoposide for 6h prior to being lifted off the plates. Single cell suspensions were made in 1X PBS 4% FBS. Cells were stained for 5 min at RT in 5 µM propidium iodide, and 25 nM Annexin V (APC, eBioscience) and were then immediately analyzed on an LSR Fortessa X20 (BD Biosciences).

*Ex vivo* stimulation and analysis of PBMCs by flow cytometry was performed essentially as described in: Lei et al.^159^. Briefly, mice were deeply anesthetized and whole blood was collected in sodium heparin tubes. Red blood cells were subjected to 2 rounds of lysis with 1X ACK lysis buffer. Leukocytes were then stimulated in RPMI+10% FBS containing LPS (1 μg/mL) in the presence of protein transport inhibitors brefeldin A and monensin for 4H. Ghost Dye 710 (Tonbo) was used as a live dead stain. Fc receptors were blocked with anti-mouse CD16/CD32 Fc shield (2.4G2, 70-0161, Tonbo), and cells were stained with antibodies against surface proteins CD3 (APC-fire, BioLegend), CD11b (APC Cy7, Tonbo), CD19 (BV605, BioLegend), Ly6C (BV650, BioLegend), MHCII (BV711, BioLegend), Ly6G (PE, Tonbo ), F4/80 (PerCP-C5.5, Tonbo). Permeabilization of cells was achieved with Foxp3/Transcription Factor Staining Buffer Kit (TNB-0607-KIT, Tonbo), and cells were stained with antibodies against intracellular TNF-α (PEC7, BioLegend). Flow cytometry was performed on a 5-laser Cytek Aurora, and Flow-Jo software was used for post-acquisition analysis.

For *ex vivo* analysis of lung cell populations on day 21 post Mtb infection, the inferior lobe was isolated and washed in 1X PBS followed by mincing and digestion for 1h in Dispase (5U/ml) at 37 °C. Following digestion tissue was passed through 70 um filters to achieve single cell suspensions. Live dead staining was performed using Ghost dye 510 (Tonbo). Fc receptors were blocked using CD16/CD32 monoclonal antibody (eBiosciences). Cells were stained with antibodies against surface proteins CD11b (BV421, BD Biosciences), CD11c (BV605, BioLegend), CD45 (BV785, BioLegend), CD170 (eFluor-488, eBiosciences), MHCII (PE, BD Biosciences), Ly6G (PerCP-Cy5.5, eBiosciences), Ly6C (APC, eBiosciences), CD206 (APCeFLuor-700 eBiosciences), B220 (APCeFluor-780, eBiosciences) . Cells were washed 2X before fixing in 4% PFA for 15 min at RT. Flow cytometry was performed on the LSR Fortessa X20, and Flow-Jo software was used for post-acquisition analysis.

### Cytoplasmic DNA enrichment

BMDMs were plated in 10 cm dishes at 1x10^7^. The next day, plates were treated with LPS and poly dA:dT as indicated. To harvest, cells were lifted with 1X PBS EDTA. Cells were washed and resuspended in 5 mL 1X PBS. Total DNA was isolated from 2% of resuspended cells and treated with 25 mM NaOH. The samples were boiled for 30 min then neutralized with 50 mM TRIS pH 8.0. The remainder of the cell suspension was pelleted at 3000 rcf for 5 min. Cells were resuspended in 500 uL cytosolic lysis buffer (50 mM HEPES pH 7.4, 150 mM NaCl, 50 µg/ mL digitonin, 10 mM EDTA) and incubated on ice for 15 min. Cells were spun down at 1000 rcf to pellet intact cells and nuclei that were then used for obtaining the membrane fraction. The supernatant was transferred to a fresh tube and spun down at 15000 rcf to remove additional organelle fragments and transferred to a fresh tube. Cytosolic protein was obtained by transferring 10% of supernatant to a fresh tube with 4x sample buffer + DTT and boiled for 5 min. Cytosolic DNA was isolated from the remaining supernatant by mixing an equal volume of 25:24:1 phenol: chloroform: isoamyl alcohol, vigorously shaken and centrifuged for 10 min at ∼21130 rcf (max speed). The aqueous phase was transferred to a fresh tube and DNA was precipitated by mixing with 300 mM sodium acetate 10 mM MgCl2 and 1 µL glycogen and 3 volumes of 100% ethanol. The DNA was pelleted by centrifugation at max speed for 20 min at 4°C. The pellet is washed with 1 mL of cold 70% ethanol and centrifuged for 5 min at max speed. The supernatant was removed, and the pellet was air dried for about 10 minutes. The pellet is resuspended with 200 µL EB. For the mitochondrial membrane fraction, the pellet of intact cells previously collected was resuspended in 500 µL membrane lysis buffer (50 mM HEPES pH 7.4, 150 mM NaCl, 1% NP-40), vortexed then centrifuged for 3 min at 7000 rcf. 50 µL of the cleared lysate is transferred to a fresh tube and received 4x sample buffer + DTT. Western blot analysis was used to check for contaminating mitochondrial proteins in the cytosolic fraction compared to the membrane fraction. qPCR was performed using total DNA diluted 1:100 and cytosolic DNA diluted 1:2 and measured *Tert*, *16s*, *ND4* and *DLoop*. The total and cytosolic reactions were normalized to *Tert* to control for variation in cell numbers.

### Immunofluorescence microscopy

MEFs were seeded at 1x10^5^ cells/well on glass coverslips in 24-well dishes. BMDMs were seeded at 3x10^5^ cells/well on glass coverslips in 24-well dishes. Cells were fixed in 4% PFA for 10 min at RT and then washed three times with PBS. Coverslips were incubated in primary antibody diluted in PBS + 5% non-fat milk + 0.1% Triton-X (PBS-MT) for 3h. Primary antibodies used in this study were phospho-MLKL (Ser358) (D6H3V) (Cell Signaling Technology, #91689; 1:200), GSDMD [EPR20859] (Abcam, ab219800, 1:100), and Tom20 (clone 2F8.1; Millipore Sigma, MABT166, 1:100).Cells were then washed three times in PBS and incubated in secondary antibodies (goat anti-rabbit Alexa Fluor 488; Invitrogen, 1:500, or goat anti-rabbit Alexa Fluor 488 and goat anti-mouse Alexa Fluor 594; Invitrogen, 1:1000) and DAPI (1:10,000) diluted in PBS-MT for 1h. Coverslips were washed twice with PBS and twice with deionized water and mounted on glass slides using Prolong Gold Antifade Reagent (Invitrogen). Z-stack images were obtained using an Olympus IX83 inverted confocal microscope equipped with 60X oil immersion objective. Quantifications were performed using Fiji ImageJ. Images were opened as separate channels and z-stacks were converted to maximum intensity projections. Each channel was thresholded such that the region of interest was masked, and then particles (nuclei) counted and the region of interest (gasdermin D) measured.

### Mitochodnrial isolation

Prior to downstream analysis, mitochondria were isolated using a Mitochondria/Cytosol Fractionation Kit (ab65320). Briefly, 5x 10^6^ BMDMs were lifted on 10 cm plates using 1X PBS-EDTA and pelleted at 600rcf for 5 minutes at 4 °C. The cells were resuspended in 500 µL 1X Cytosolic Extraction Buffer Mix provided by the kit without DTT or Protease Inhibitors and incubated on ice for 10 min. Cells were homogenized on ice with 100 passes using the tight-fitting pestle B to lyse the cells followed by differential centrifugation. Homogenate was centrifuged at 700 rcf for 10 min at 4 °C to clear un-lysed cells and nuclei. The supernatant was collected and placed into a fresh tube and centrifuged at 10,000 rcf for 30 min at 4 °C. The supernatant was collected into a fresh tube and 4X sample buffer with 5 mM DTT was added. The pellet was either resuspended in 1X PBS for flow cytometry or lysed in sample buffer with 5 mM DTT for western analysis.

### Protein quantification by immunoblot

Cells were washed with PBS and lysed in 1X RIPA buffer with protease and phosphatase inhibitors, with the addition of 1 U/ml Benzonase to degrade genomic DNA. Proteins were separated by SDS-PAGE and transferred to nitrocellulose membranes. Membranes were blocked for 1h at RT in LiCOR Odyssey blocking buffer or TBS with 5% BSA. Blots were incubated overnight at RT with the following antibodies: Beta Actin (Abcam, 1:2000), DRP1 (Cell Signaling, 1:1000); pDRP1 Ser616 (Cell Signaling, 1:1000), pDRP1 Ser637 (Cell Signaling, 1:1000),TFAM (Millipore, 1:1000), VDAC (Protein Tech, 1:1000). Membranes were incubated with appropriate secondary antibodies for 2h at RT prior to imaging on a LiCOR Odyssey Fc Dual-Mode Imaging System.

### GSDMD/CASP11 protein expression and purification

The cDNAs encoding human and mouse GSDMD were inserted into a modified pET28(a) vector with an N-terminal Avi-His6-SUMO-tag. The proteins were expressed in *E. coli* BL21 in regular LB medium (BD). When OD600 reached 1.0, the cells were induced with 0.4 mM isopropyl β-D-1-thiogalactopyranoside (IPTG) and cultured overnight at 16 °C. The *E. coli* cells were harvested by centrifugation at 5000 rpm for 10 min. Then, the cells were lysed in a lysis buffer containing 50 mM Tris, 300 mM NaCl, pH 8.0 by sonication on ice. After centrifugation at 4 °C, 16000 rpm for 30 min, the supernatants containing the target proteins were loaded onto a nickel-NTA column equilibrated with the lysis buffer. After washing the column with 200 ml washing buffer (20 mM Tris, 500 mM NaCl, 25 mM Imidazole, pH 7.5), the target proteins were eluted with the elution buffer (20 mM Tris, 150 mM NaCl, 250 mM Imidazole, pH 7.5). The Avi-His6-SUMO tag was cleaved using SUMO protease at 4 °C overnight and removed using a nickel-NTA column. Human and mouse GSDMD were further purified by gel filtration chromatography using a Superdex200 (16/60 GL) column (GE Healthcare) in a running buffer containing 20 mM Tris, 150 mM NaCl, at pH 7.5.

To express inactive mouse caspase-11 catalytic domain (residue 100 to 373), it was cloned into a modified pET28(a) vector with an N-terminal His6-SUMO-tag. Cys254 in the active site of caspase-11 was mutated to alanine. Asp285 was replaced with a thrombin cleavage site with the sequence LVPRGS. The inactive mouse caspase-11 catalytic domain was expressed and purified using the same protocol used for GSDMD. After purification on a Superdex200 column, the inactive mouse caspase-11 catalytic domain was incubated with thrombin at 4 °C overnight and further purified on a Superdex200 column in the same running buffer as for GSDMD. To generate active mouse caspase-11 catalytic domain (residue 100 to 373), it was cloned into a modified pET28(a) vector with an N-terminal His6-SUMO-tag and a C-terminal His6-tag. There is a thrombin recognition site LVPRGS between His6- and SUMO-tag. Asp285 was replaced with a thrombin cleavage site. The His6-SUMO-mouse caspase-11 catalytic domain-His6 protein was expressed in *E.coli* BL21 and purified by nickel-NTA column using the same protocol used for GSDMD. The eluted protein from nickel-NTA column was incubated with SUMO protease and thrombin at 4 °C overnight. The active mouse caspase-11 catalytic domain-His6 protein was purified by a nickel-NTA column and SUMO-tag was removed. Then, it was further purified on a Superdex200 column in the same running buffer as for GSDMD.

All the proteins were stored in a buffer containing 20 mM Tris, 150 mM NaCl, 5 mM DTT, at pH 7.5. To test if the active mouse caspase-11 catalytic domain protein can cleave GSDMD or not, GSDMD was diluted into the reaction buffer containing 50 mM HEPES, 3 mM EDTA, 150 mM NaCl, 0.005% (v/v) Tween-20, 10 mM DTT, pH 7.5. The final concentration of GSDMD was 0.5 mg/ml and the ratio of GSDMD and caspase-11 was 10 : 1 (w/w). After incubation at 37 °C for 1 h, the samples were analyzed by SDS-PAGE.

### In vivo Mtb infections

All infections were performed using procedures approved by Texas A&M University Institutional Care and Use Committee. The Mtb inoculum was prepared as described above. Age- and sex-matched mice were infected via inhalation exposure using a Madison chamber (Glas-Col) calibrated to introduce 100-200 CFUs per mouse. For each infection, approximately 5 mice were euthanized immediately, and their lungs were homogenized and plated to verify an accurate inoculum. Infected mice were housed under BSL3 containment and monitored daily by lab members and veterinary staff. At the indicated time points, mice were euthanized, and tissue samples were collected. Blood was collected in serum collection tubes, allowed to clot for 1-2 h at room temperature, and spun to separate serum. Serum cytokine analysis was performed by Eve Technologies. Organs were divided to maximize infection readouts (CFUs: left lobe lung and ½ spleen; histology: 2 right lung lobes and ¼ spleen; RNA: 1 right lung lobe and ¼ spleen). For histological analysis organs were fixed for 24 h in neutral buffered formalin and moved to ethanol (lung, spleen). Organs were further processed as described below. For cytokine transcript analysis, organs were homogenized in Trizol Reagent, and RNA was isolated as described below. For CFU enumeration, organs were homogenized in 5 ml PBS + 0.1% Tween-80, and serial dilutions were plated on 7H10 plates. Colonies were counted after plates were incubated at 37°C for 3 weeks.

### Histopathology

Lungs and spleens fixed with 10% neutral buffered formalin followed by 70% ethanol were subjected to routine processing, embedded in paraffin, and 5 μm sections were cut and stained with hematoxylin and eosin (H&E) or acid-fast stain (Diagnostic BioSystems). A board certified veterinary pathologist performed a masked evaluation of lung sections for inflammation using a scoring system: score 0, none; score 1, up to 25% of fields; score 2, 26-50% of fields; score 3, 51-75% of fields; score 4, 76-100% of fields. To quantify the percentage of lung fields occupied by inflammatory nodules, scanned images of at least 2 sections of each lung were analyzed using Fiji ImageJ (NIH) to determine the total cross-sectional area of inflammatory nodules per total lung cross sectional area^160^.

### Immuno-multiplex assay

Sera was analyzed by Eve Technologies: Mouse Cytokine Array/Chemokine Array 13-plex Secondary Panel (MD13). Briefly, sera was isolated following decapitation in Microtainer serum separator tubes (BD Biosciences) followed by 2x sterile filtration with Ultrafree-MC sterile filters, 10 min at 10,000 rpm (Millipore Sigma). For analysis sera was prediluted 1:1 to a final volume of 100μl in 1x PBS pH 7.4 and assayed/analyzed in duplicate.

### Drosophila husbandry and strains

The following strains were obtained from Bloomington Drosophila Stock Center: w1118 and DaGal4. UAS-hLRRK2-G2019S and UAS-hLRRK2-G2019S-K1906M were kindly provided by C. Elliott and W. Smith. CGGal4 was kindly provided by C. Thummel.

All flies were reared on standard yeast and cornmeal-based diet at 25°C and 65% humidity on a 12 hr light/dark cycle, unless otherwise indicated. The standard lab diet (cornmeal-based) was made with the following protocol: 14g Agar/165.4g Malt Extract/ 41.4g Dry yeast/ 78.2g Cornmeal/ 4.7ml propionic acid/ 3g Methyl 4-Hydroxybenzoate/ 1.5L water. In order to standardize metabolic results, fifty virgins were crossed to 10 males and kept in bottles for 2-3 days to lay enough eggs. Wet folded filters (GE healthcare, CAT No.10311843) were inserted in bottles after parental flies were removed. Progeny of crosses was collected for 3–4 days after initial eclosion. Collected progeny were then transferred to new bottles to allow them mate for 2 days (representing unique populations). All these flies were reared on a standard lab diet at 25°C and 65% humidity on a 12 hr light/dark cycle, unless otherwise indicated. Around 20 female flies were then separated into each vial (before mock or oral infection treatment) for 10 days at 25°C and 65% humidity on a 12 hr light/dark cycle. Post-mated Female flies were used for all experiments due to sensitivity to P.e. infection. The UAS-hLRRK2-G2019S, UAS-hLRRK2-G2019S-K1906M, CGGal4, and DaGal4 transgenic lines were backcrossed 10x into the w1118 background that was used as a control strain, with continued backcrossing every 6-8 months to maintain isogeneity.

### Oral *P. entomophila* fly infection

Bacterial entomopathogen strain *Pseudomonas entomophila* (P.e) was grown in LB medium at 29°C, shaking at 200 rpm overnight. Fresh bacterial cultures were generated daily. The liquid cultures were poured into a sterile centrifuge flask and centrifuged at 4000 g at 4°C for 15 minutes. The liquid LB medium was removed from the centrifuge flask, and the bacterial pellet was resuspended in a small amount of LB medium. The final bacteria concentration (of the resuspended pellet, OD600=50-60 for P.e) was adjusted by diluting with additional LB medium. Bacterial cultures were routinely genotyped for accuracy and reproducibility of experiments.

Next, 2.5% and 5% sucrose (in sterile water) were prepared fresh. The 5% sucrose solution was mixed with an equal volume of bacteria solution (at OD600=50-60 for P.e) to create the solution used for oral infection (feeding). NOTE: This OD were used for all experiments except for survival analysis. 10 day-old mated female flies (20 per vial) were transferred into a fly food vial containing a filter paper that totally covers the food and was soaked with a solution consisting of 185 μL either bacterial oral infection mix (for infections) or 2.5% sucrose, for unchallenged (mock) controls. Drosophila were always infected at 3:00 - 4:00pm to ensure diurnal reproducibility, and subsequently incubated at 25°C and 65% humidity on a 12 hours light/dark cycle for required infection times (16-20 hours for P.e) while feeding.

### *Drosophila* immunostaining, Nile Red staining, and microscopy

For fat body/adipose immunostaining, carcass was dissected (with all of the eggs and intact intestines removed) in PBS and fixed with 4% paraformaldehyde for 20 min at room temperature, washed 3 times with PBS containing 0.2% Triton X-100 (PBST) and then block in blocking buffer (5% BSA in PBST) for 1 h. Primary antibodies; anti-ATP5A from Abcam (ab14748, 1:500) was applied overnight at 4°C. Samples were then incubated with Alexa Flour-conjugated secondary antibodies (Jackson Immunoresearch, 1:500), and fresh Nile Red solution (1µl of 0.004% Nile Red Solution in 500 µl PBS) with Hoechst (DAPI; 1:500), overnight at 4°C, followed by rinsing with PBS. Confocal images were collected using a Nikon Eclipse Ti confocal system (utilizing a single focal plane) and processed using the Nikon software and Adobe Photoshop.

### Survival analysis

Flies were infected with P.e at OD600=30-40 as described above and kept at 25°C for 16 hours. After overnight feeding (flies were always orally infected at 3:00 - 4:00pm), bacterial infected flies were transferred from bacterial infection vials to vials containing standard lab food. Flies were transferred every day to a fresh vial for the first two days, and every two days after, and dead flies were counted (and removed) when changing vials.

### Quantification and statistical analysis

All data are representative of two or more independent experiments with n=3 or greater. For all quantifications, n represents the number of biological replicates, and error bar represents SEM. Statistical significance was determined using either the unpaired t test, one-way ANOVA with Tukey post hoc test, or two-way ANOVA with Tukey post hoc test where multiple comparisons were necessary, in GraphPad Prism Software, and expressed as P values. For mouse experiments, we estimated that detecting a significant effect requires two samples to differ in CFUs by 0.7e^10. Using a standard deviation of 0.35e^10 for each population, we calculated that a minimum size of 5 age- and sex-matched mice per group per time point is necessary to detect a statistically significant difference by a t-test with alpha (2-sided) set at 0.05 and a power of 80%. Therefore, we used a minimum of 5 mice per genotype per time point to assess infection-related readouts. For statistical comparison, each experimental group was tested for normal distribution. Data were tested using a Mann-Whitney test.

