## Supplemental Figures for "Mitochondrial dysfunction promotes alternative gasdermin D-mediated inflammatory cell death and susceptibility to infection"

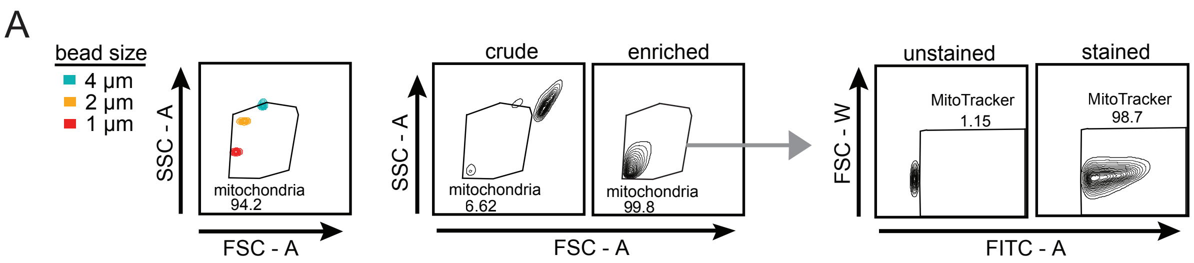
A
bead size
4 µm
2 µm
1 µm
SSC - A
FSC - A
mitochondria
94.2
crude
enriched
SSC - A
mitochondria
6.62
mitochondria
99.8
FSC - A
unstained
stained
FSC - W
MitoTracker
1.15
MitoTracker
98.7
FITC - A

Figure S1

**Figure S1**

**A.** Gating strategy used to identify pure populations of mitochondria during flow cytometry analysis. (Left plot) Bead size standards defining gate. (middle plot) Crude and pure mitochondrial populations within the size exclusion gate. (right plot) Staining with 100 nM MitoTracker green to validate purity of mitochondria.

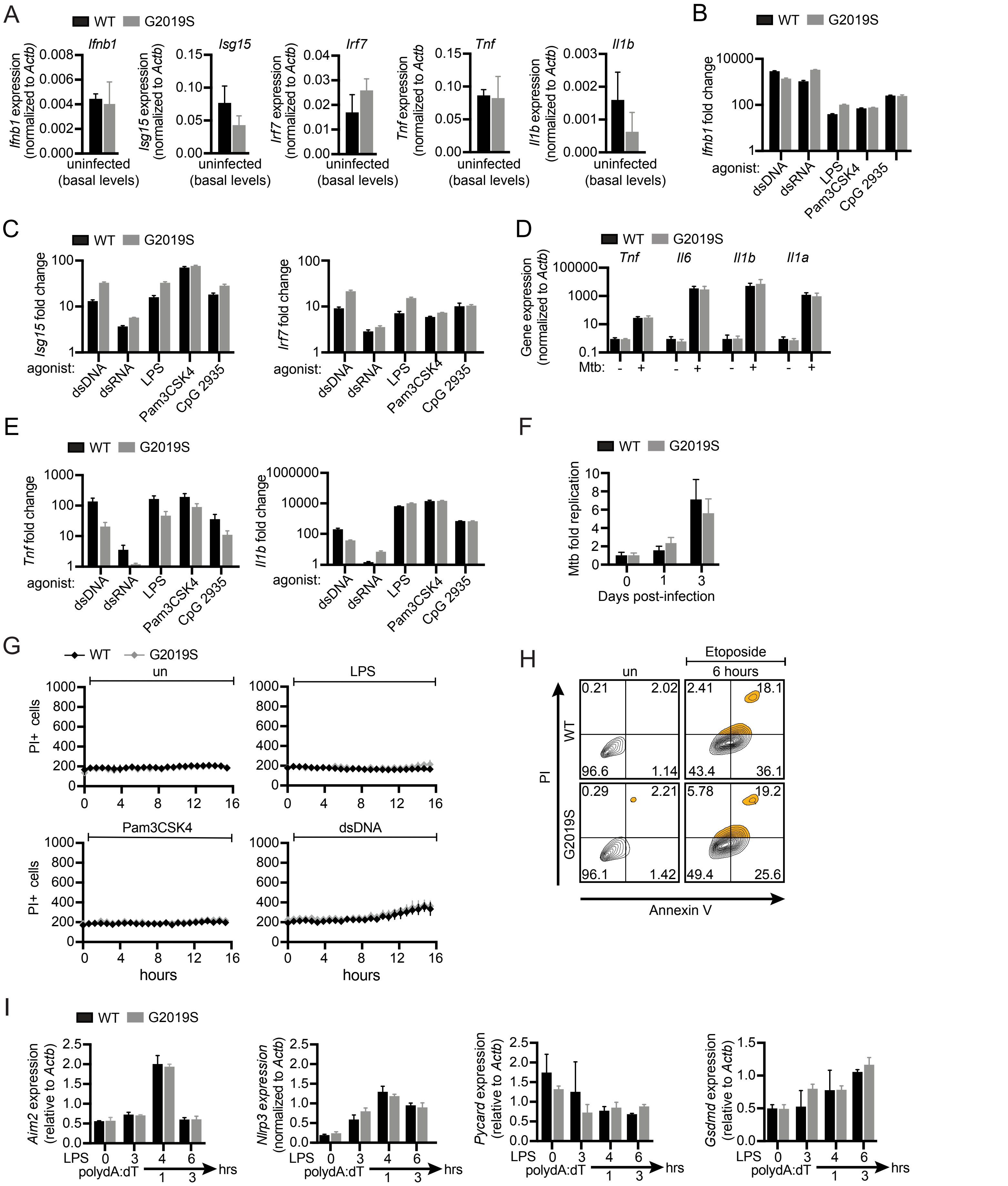
A
WT
G2019S
Ifnb1
Ifnb1 expression (normalized to Actb)
Isg15
Isg15 expression (normalized to Actb)
Irf7
Irf7 expression (normalized to Actb)
Tnf
Tnf expression (normalized to Actb)
Il1b
Il1b expression (normalized to Actb)
uninfected (basal levels)
B
Ifnb1 fold change
agonist:
dsDNA
dsRNA
LPS
Pam3CSK4
CpG 2935
C
Isg15 fold change
Irf7 fold change
D
Gene expression (normalized to Actb)
Il6
Il1a
Mtb:
E
Tnf fold change
Il1b fold change
F
Mtb fold replication
Days post-infection
G
PI+ cells
un
hours
H
Etoposide
6 hours
PI
Annexin V
I
Aim2 expression (relative to Actb)
Nlrp3 expression (normalized to Actb)
Pycard expression (relative to Actb)
Gsdmd expression (relative to Actb)
LPS
polydA:dT
hrs

Figure S2

**Figure S2**

**A.** Basal transcripts of *Ifnb1*, *Isg15*, *Irf7*, *Tnf*, and *Il1b* in wild-type and *Lrrk2*$^{G2019S}$ BMDMs as measured by RT-qPCR. **B.** *Ifnb1* fold change at 4h post-treatment with a panel of agonists: dsDNA (ISD) 1 µg/ml, dsRNA (Poly I:C) 1 µg/ml, LPS 100 ng/ml, Pam3CSK4 100 ng/ml, CpG 2935 1 µM in wild-type and *Lrrk2*$^{G2019S}$ BMDMs. **C.** As in B, but for interferon stimulated genes (*Isg15* and *Irf7*). **D.** *Tnf*, *Il6*, *Il1b* and *Il1a* gene expression as measured by qRT-PCR 4h post-Mtb infection (MOI 10) in wild-type and *Lrrk2*$^{G2019S}$ BMDMs. **E.** As in B, but for *Tnf* and *Il1b.* **F.** CFU quantification of Mtb replication in wild-type and *Lrrk2*$^{G2019S}$ BMDMs over 3 days (MOI 1). **G.** PI staining over a time course in untreated wild-type and *Lrrk2*$^{G2019S}$ BMDMs or in the presence of LPS 100 ng/ml, Pam3CSK4 100 ng/ml, dsDNA (ISD) 1 µg/ml. **H.** Flow cytometry of PI staining (y-axis) and annexin-V staining (x-axis) in WT and G2019S BMDMs treated with or without etoposide 50 µM for 6 hrs. **I.** *Aim2*, *Nlrp3*, Pycard, and *Gsdmd* transcript levels in wild-type and *Lrrk2*$^{G2019S}$ BMDMs as measured by RT-qPCR following AIM2 inflammasome stimulation or LPS priming at indicated times.

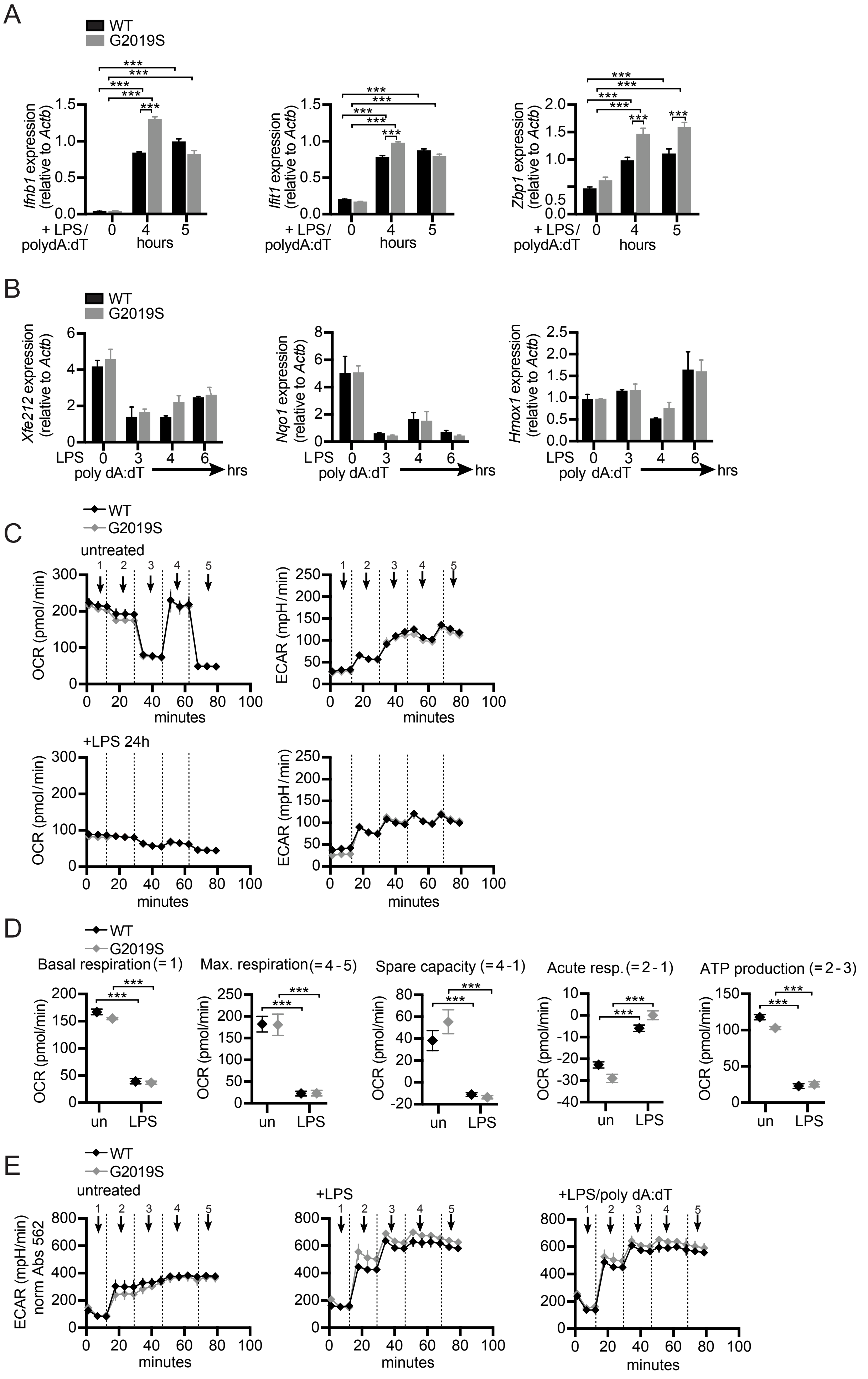
A
WT
G2019S
Ifnb1 expression (relative to Actb)
Ifit1 expression (relative to Actb)
Zbp1 expression (relative to Actb)
+ LPS/ polydA:dT
hours
B
Xfe212 expression (relative to Actb)
Nqo1 expression (relative to Actb)
Hmox1 expression (relative to Actb)
LPS
poly dA:dT
hrs
C
untreated
+LPS 24h
OCR (pmol/min)
ECAR (mpH/min)
minutes
D
Basal respiration (= 1)
Max. respiration (= 4 - 5)
Spare capacity (= 4 -1)
Acute resp. (= 2 - 1)
ATP production (= 2 - 3)
un
LPS
E
untreated
+LPS
+LPS/poly dA:dT
ECAR (mpH/min) norm Abs 562

Figure S3

**Figure S3**

**A.** Total transcript levels of *Ifnb1* and ISGs after AIM2 stimulation in wild-type and *Lrrk2*$^{G2019S}$ measured by RT-qPCR. **B.** As in A but for components of the NRF2 regulon. **C.** Oxygen consumption rate (OCR) measured by Agilent Seahorse Metabolic Analyzer for wild-type and *Lrrk2*$^{G2019S}$ BMDMs +10 ng/ml LPS 24h. **D.** Spare respiratory capacity, maximal respiration, acute response, and non-mitochondrial oxygen consumption of untreated and LPS-treated (24h) wild-type and *Lrrk2*$^{G2019S}$ BMDMs. **E.** Extracellular acidification rate (ECAR) measured by Agilent Seahorse Metabolic Analyzer for wild-type and *Lrrk2*$^{G2019S}$ BMDMs +10 ng/ml LPS 3h, or AIM2 stimulation for 3h.

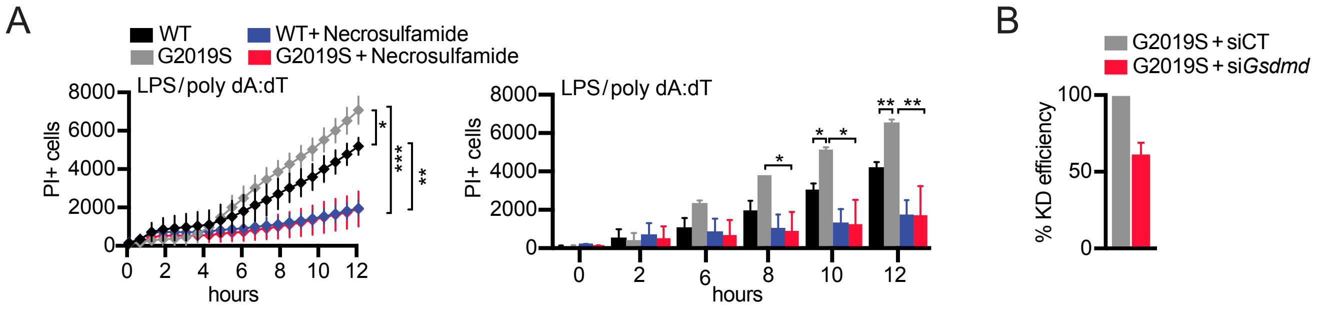
A
WT
G2019S
WT+Necrosulfamide
G2019S+Necrosulfamide
LPS/poly dA:dT
PI+ cells
hours
B
G2019S+siCT
G2019S+siGsdmd
% KD efficiency

Figure S4

**Figure S4.**

**A.** PI staining over a time course of AIM2 activation in wild-type and *Lrrk2*$^{G2019S}$ BMDMs +/- necrosulfamide treatment (20 μM). Quantification of PI incorporation at each hour on the right. **B.** *Gsdmd* knockdown efficiency as measured by RT-qPCR.

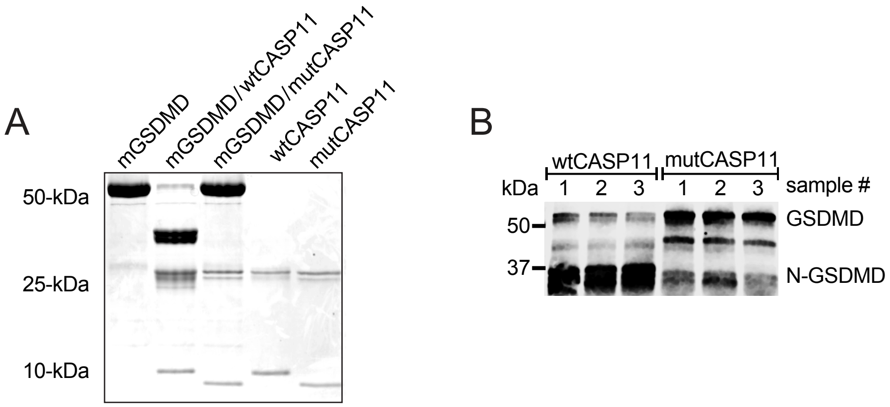
A
mGSDMD
mGSDMD/wtCASP11
mGSDMD/mutCASP11
wtCASP11
mutCASP11
50-kDa
25-kDa
10-kDa
B
wtCASP11
mutCASP11
kDa
1
2
3
1
2
3
sample #
50
37
GSDMD
N-GSDMD

Figure S5

**Figure S5.**

**A.** Silver stain of full length GSDMD, N-GSDMD cleaved by the catalytic domain of CASP11, full length GSDMD uncleaved by the mutant catalytic domain of CASP11, active CASP11 (wtCASP11) and mutant C254A CASP11 (mutCASP11). **B.** Western blot analysis of mitochondria isolated from BMDMs cleaved with either wtCASP11 or uncleaved by mutCASP11.

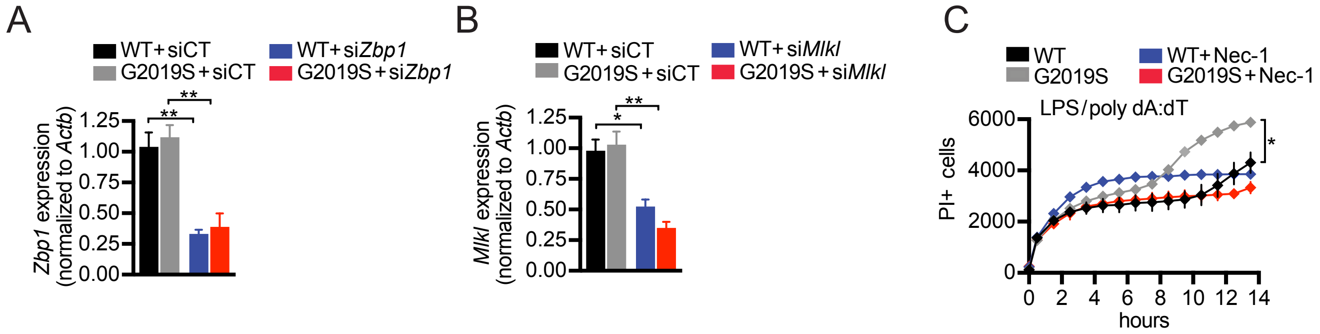
A
WT+siCT
WT+siZbp1
G2019S+siCT
G2019S+siZbp1
Zbp1 expression (normalized to Actb)
1.25
1.00
0.75
0.50
0.25
0.00
**
**
B
WT+siCT
WT+siMlkl
G2019S+siCT
G2019S+siMlkl
Mlkl expression (normalized to Actb)
1.25
1.00
0.75
0.50
0.25
0.00
*
**
C
WT
WT+Nec-1
G2019S
G2019S+Nec-1
LPS/poly dA:dT
PI+ cells
6000
4000
2000
0
0 2 4 6 8 10 12 14
hours
*

Figure S6

**Figure S6.**

**A.** *Zbp1* knockdown efficiency as measured by RT-qPCR. **B.** As in A but for *Mlkl.* **C.** PI staining over a time course of AIM2 activation in wild-type and *Lrrk2*$^{G2019S}$ BMDMs +/- the RIPK1 inhibitor necrostatin-1 (10 μM).

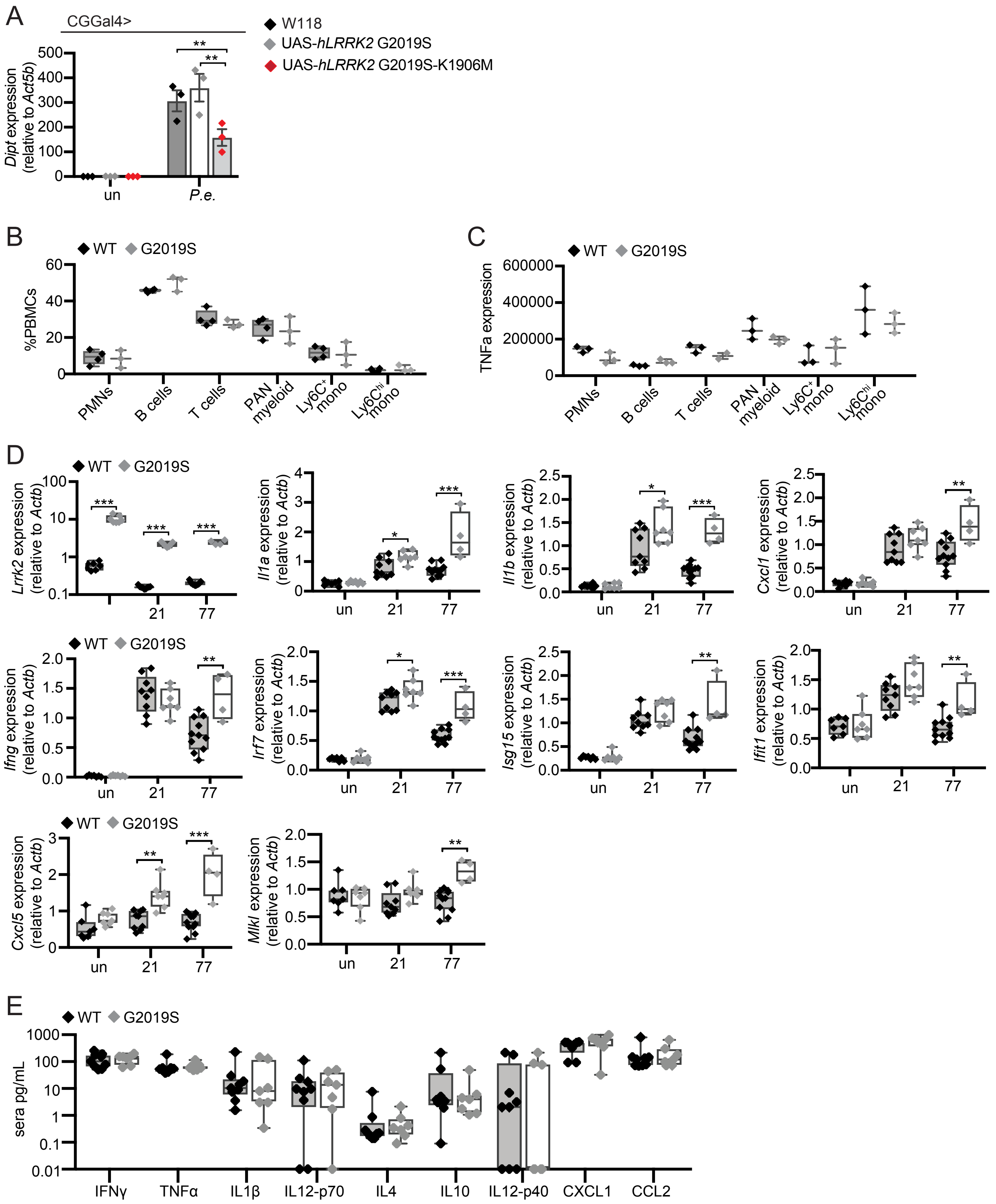
A
CGGal4>
W118
UAS-hLRRK2 G2019S
UAS-hLRRK2 G2019S-K1906M
Dipt expression (relative to Act5b)
un
P.e.
B
WT
G2019S
%PBMCs
PMNs
B cells
T cells
PAN myeloid
Ly6C+ mono
Ly6Chi mono
C
TNFa expression
D
Lrrk2 expression (relative to Actb)
Il1a expression (relative to Actb)
Il1b expression (relative to Actb)
Cxcl1 expression (relative to Actb)
Ifng expression (relative to Actb)
Irf7 expression (relative to Actb)
Isg15 expression (relative to Actb)
Ifit1 expression (relative to Actb)
Cxcl5 expression (relative to Actb)
Mlkl expression (relative to Actb)
un
21
77
E
sera pg/mL
IFNγ
TNFα
IL1β
IL12-p70
IL4
IL10
IL12-p40
CXCL1
CCL2

Figure S7

**Figure S7.**

**A.** RT-qPCR of the inflammatory mediators *Dipt* in wild-type (W118), hLRRK2-, hLRRK2-G2019S-, and hLRRK2 G2019S-K1906M-expressing flies after *P.e.* infection (20 hours). Gene expression is shown relative to an *Act5c* control. **B.** PBMC cell populations in resting cells from wild-type and *Lrrk2*$^{G2019S}$ mice as measured by flow cytometry. **C.** TNF-α protein expression as measured by MFI in resting PBMC cell populations. **D.** RT-qPCR of *Lrrk2* and inflammatory cytokines from total RNA recovered from lung homogenates from uninfected and Mtb-infected wild-type and *Lrrk2*$^{G2019S}$ mice at day 21 and 77 post-infection. **E.** Serum cytokines in wild-type and *Lrrk2*$^{G2019S}$ mice measured by cytokine array at day 21 after Mtb infection.
